# Thymosin β4 mediates vascular protection via regulation of Low Density Lipoprotein Related Protein 1 (LRP1)

**DOI:** 10.1101/535351

**Authors:** Sonali Munshaw, Susann Bruche, Jyoti Patel, Andia Redpath, Karina N. Dubé, Rachel Davies, Giles Neal, Regent Lee, Ashok Handa, Keith M. Channon, Nicola Smart

**Affiliations:** Burdon Sanderson Cardiac Science Centre, Department of Physiology, Anatomy & Genetics, University of Oxford, Sherrington Building, South Parks Road, Oxford OX1 3PT, UK; BHF Centre of Research Excellence, Division of Cardiovascular Medicine, John Radcliffe Hospital, University of Oxford, UK; UCL-Institute of Child Health, 30 Guilford Street, London WC1N 1EH, UK; Nuffield Department of Surgical Sciences, University of Oxford, Oxford, UK

## Abstract

Vascular stability and tone are maintained by contractile smooth muscle cells (VSMCs). However, injury-induced growth factors stimulate a contractile-synthetic phenotypic switch which promotes atherosclerosis and susceptibility to abdominal aortic aneurysm (AAA). As a regulator of embryonic VSMC differentiation, we hypothesised that Thymosin β4 (Tβ4) may additionally function to maintain healthy vasculature and protect against disease throughout postnatal life. This was supported by identification of an interaction with Low density lipoprotein receptor related protein 1 (LRP1), an endocytic regulator of PDGF-BB signalling and VSMC proliferation. LRP1 variants have been identified by GWAS as major risk loci for AAA and coronary artery disease. Tβ4-null mice display aortic VSMC and elastin defects, phenocopying LRP1 mutants and suggesting compromised vascular integrity. We confirmed predisposition to disease in models of atherosclerosis and AAA. Diseased vessels and plaques were characterised by accelerated contractile-synthetic VSMC switching and augmented PDGFRβ signalling. In vitro, enhanced sensitivity to PDGF-BB, upon loss of Tβ4, coincided with dysregulated endocytosis, leading to increased recycling of LRP1-PDGFRβ and reduced lysosomal targeting. Our study identifies Tβ4 as a key regulator of LRP1 for maintaining vascular health, providing insight which may reveal useful therapeutic targets for modulation of VSMC phenotypic switching and disease progression.

## Introduction

Vascular diseases are a leading cause of morbidity and mortality worldwide. The integrity of the blood vessel wall is key to resisting the shear forces of flowing blood and preventing aneurysmal dilatation and rupture. Endothelial damage triggers macrophage infiltration and lipid accumulation, leading to atherosclerotic plaque formation, compromised vascular stability and susceptibility to aortic aneurysm. Genome-wide association studies (GWAS) have provided important insights into genetic predisposition for aortic disease, however, a major challenge in the ‘post-genomic era’ remains to elucidate the molecular mechanisms through which GWAS ‘hits’ influence disease phenotype(1, 2).

Vascular stability is determined by competing degenerative (macrophage-secreted proteases, elastolysis and smooth muscle cell (VSMC) apoptosis) and regenerative mechanisms (VSMC survival and synthesis of an elastin-rich extracellular matrix (ECM)). VSMCs, in their contractile state, play an essential role in regulating vascular tone. In disease, however, growth factors such as platelet-derived growth factor (PDGF)-BB, secreted by damaged endothelium and immune cells, induce a contractile-synthetic phenotypic switch in VSMCs to facilitate proliferation, migration and ECM synthesis. Although reparative in the short term, chronic VSMC dedifferentiation leads to vascular thickening and stiffness, exacerbates inflammation and promotes atherosclerotic lesion development(3). Indeed, inhibiting VSMC phenotypic transformation has been shown to attenuate progression of vascular disease(4).

Understanding mechanisms of embryonic VSMC differentiation may inform strategies for maintaining contractile phenotype to prevent the pathological changes that underlie vascular disease. *TMSB4X*, encoding the actin monomer (G-actin) binding protein, is the most abundant transcript in healthy and abdominal aortic aneurysm (AAA) patient aorta(5), yet its endogenous roles in the adult vasculature have not been explored(6). We previously defined an essential requirement for Tβ4 in mural cell differentiation in the developing mouse embryo(7). A proportion of Tβ4-null embryos die at E12.5 with vascular haemorrhage, coincident with a reduction in VSMC coverage of their developing vessels(7). These findings are consistent with similar roles for Tβ4 in smooth muscle differentiation in the coronary(8) and yolk sac vasculature(9). Despite the severe embryonic defects, the majority of Tβ4KO mice survive to adulthood(7). To determine whether Tβ4 is required to maintain vascular integrity postnatally, we sought to investigate the phenotype of adult vessels. Tβ4 null aortas were significantly dilated with highly disorganised and irregular VSMC morphology, accompanied by aberrant elastin deposition, suggesting severely compromised vascular stability. Such medial layer defects are observed in patients with AAA(10) and atherosclerosis(11). In keeping with this, we confirmed predisposition of global and VSMC- specific Tβ4 loss-of-function mice to disease in experimental models of aortic aneurysm (1mg/kg/day Angiotensin II) and atherosclerosis (ApoE-/-hypercholesterolaemia). Aortic dilatation and unstable atherosclerotic plaques in mutant mice were characterised by enhanced contractile-synthetic VSMC switching and proliferation, underpinned by dysregulated PDGFRβ signalling.

Seeking insight into the underlying mechanisms, we identified a putative interaction between Tβ4 and Low density lipoprotein (LDL) receptor related protein 1 (LRP1), which functions in VSMC development and protection, in part, by regulating growth factor signalling. Notably, LRP1 variants have been identified by GWAS as major risk loci for AAA(12), carotid(13) and coronary artery disease(14). LRP1 is involved in vascular remodelling, inflammation, differentiation and cell migration(15), roles shared with Tβ4, and has been shown, in animal studies, to protect against aneurysm and atherosclerosis(16, 17). LRP1 functions as an endocytic co-receptor for PDGFRβ to both potentiate and attenuate downstream signalling activity(18). From endosomes, the LRP1-PDGFRβ complex may be recycled to the cell membrane or targeted for lysosomal degradation; such trafficking critically determines sensitivity to PDGF isoforms and cellular responses. We report the hyperactivation of LRP1-PDGFRβ signalling in Tβ4-null aortic VSMCs and find that augmented sensitivity correlates with increased recycling of LRP1-PDGFRβ to the cell surface, concomitant with reduced lysosomal targeting. Taken together, these findings suggest a requirement for Tβ4 in the normal attenuation of VSMC PDGFRβ signalling, via lysosomal targeting of the receptor, to maintain contractile VSMC phenotype and vascular stability. Our study reveals novel insight into the mechanism by which Tβ4 controls LRP1-mediated VSMC responses to protect against vascular disease. Given that LRP1 has been implicated in multiple GWAS as a key regulator of vascular protection, identification of an important molecular regulator may enable the development of novel therapies for modulation of VSMC phenotypic switching and disease progression.

### A role for Tβ4 in maintenance of adult vascular stability

Whilst vascular defects cause lethality in a proportion of *Tmsb4x*/Tβ4 null embryos (36% of Tβ4^−/Y^ males; 16% of Tβ4^−/−^ females)(7), most survive to adulthood. This prompted us to investigate whether vessel structure and function were entirely normal in viable adults. Due to the higher mortality in male embryos, we focussed our studies primarily on male mice. Compared with control Tβ4^+/Y^ aortas, Tβ4^−/Y^ aortas of 12-16 week old mice were significantly dilated (mean 1.5-fold, determined histologically from both abdominal (AA) and thoracic (TA) sections; Figure 1, A-B) and a 1.4-fold elevation of systemic vascular compliance (stroke volume/pulse pressure) was revealed by combined MRI and arterial blood pressure measurements (Figure 1C). Moreover, aberrant elastin lamellar integrity suggested severely compromised vascular stability (3.6-fold more elastin breaks per section and higher elastin damage score, systematically assessed according to a previously defined scoring system(19) and examples in Supplemental Figure 1; Figure 1, D-F). Masson’s Trichrome staining revealed limited aortic fibrosis at 13 weeks, although, by 35 weeks, Tβ4^−/Y^ aortas displayed more collagen within the medial layer, compared with Tβ4^+/Y^ control aortas (Figure 1G). Due to the requirement for Tβ4 in VSMC differentiation ((7, 8, 20)), we assessed the phenotype of medial layer VSMCs by measuring expression of established contractile (*Sm22a*, αSMA) and synthetic markers (*Tropomysin*, L-caldesmon)(21), by quantitative reverse transcription real-time PCR (qRT-PCR) and immunofluorescence (Figure 1 H-J). Significant reductions (<0.5 fold) in the ratio of contractile/synthetic protein expression confirmed an overall shift towards more synthetic VSMCs in Tβ4^−/Y^ aortas, which was consistent with the disorganised VSMC morphology observed (Figure 1J). While Tβ4^+/Y^ aortas contained regularly aligned, elongated VSMCs, Tβ4^−/Y^ aortas frequently contained clusters of small, densely packed, cuboidal cells. Collectively, these data suggest that Tβ4^−/Y^ VSMCs are less differentiated and more synthetic in phenotype than Tβ4^+/Y^ VSMCs and that these defects may compromise vascular stability and predispose to disease.

**Figure 1.**
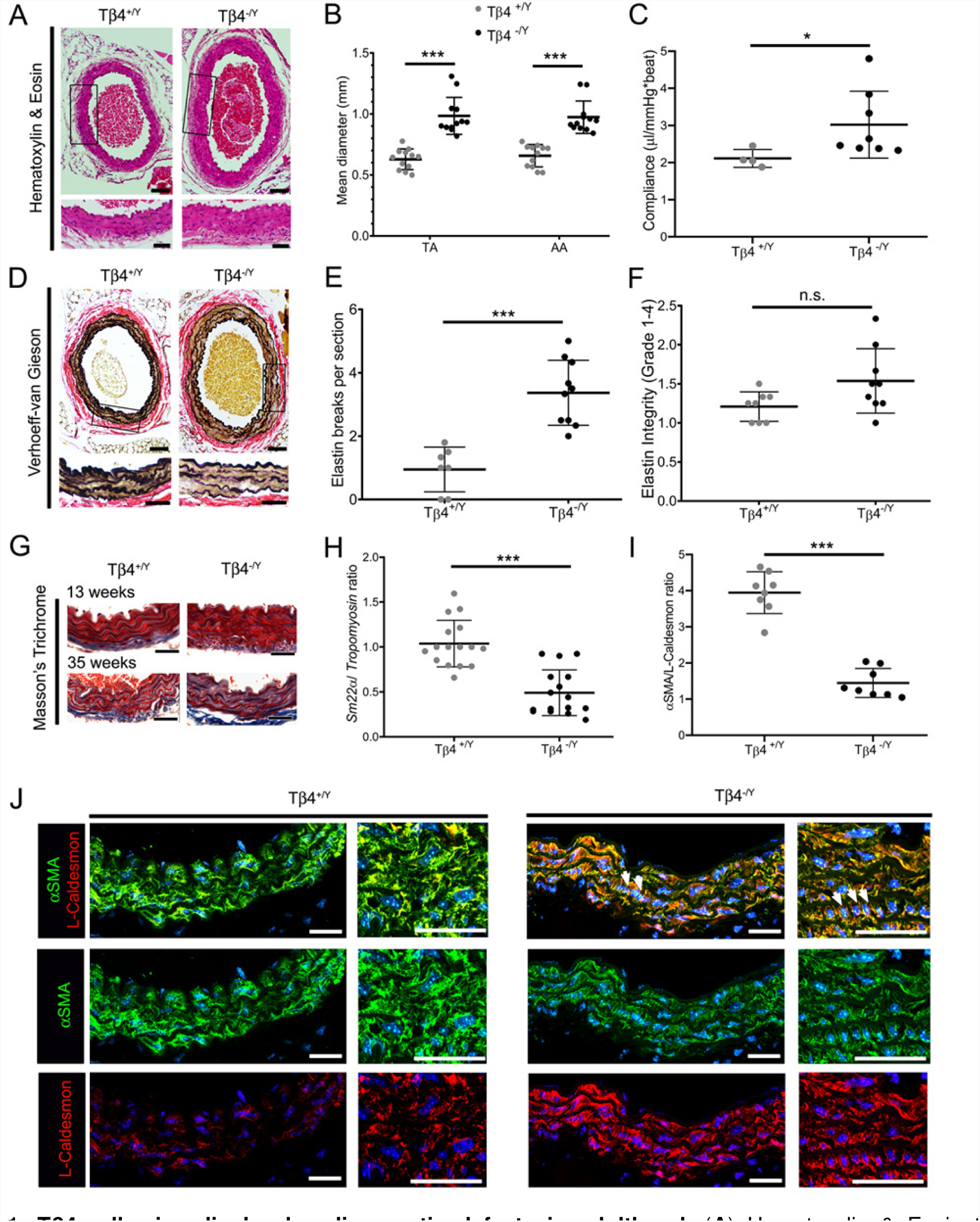
Tβ4-null mice display baseline aortic defects in adulthood. (**A**) Hematoxylin & Eosin to assess morphology and abdominal aortic dilatation, quantified in **B**. MRI and arterial blood pressure measurements of vascular compliance (stroke volume/pulse pressure; **C**). Verhoeff-van Gieson staining (**D**) assessed elastin integrity, quantified both by number of breaks per section (**E**) and by an elastin damage score (**F**; 1-4 = low-high); mean of 6 sections per aorta. Masson’s Trichrome staining (**G**) to visualise fibrosis in young (13 week old) and aging (35 week old) adult mice. Contractile/synthetic smooth muscle markers were assessed at the mRNA (qRT-PCR; **H**) and protein levels (Immunofluorescence; **I**, **J**). Altered marker profile was accompanied by altered morphology, with dense clusters of cells with a cuboidal (synthetic), rather than elongated (contractile), appearance (white arrowheads). Data are presented as mean ± SD, with each data point representing an individual animal. Significant differences were calculated using Mann Witney non-parametric tests (**B, C, E, F, I**) or a two-tailed unpaired Student’s t-test (**H**). n.s. = not significant; *p ≤ 0.05; ***: p≤0.001 for Tβ4^+/Y^ vs Tβ4^−/Y^. Scale bars: **A**, **D** and **G**: 100μm (whole); 50μm (zoom); **J**: all 50μm.

Perturbations in embryonic vascular development can predispose to aortic disease later in life(22) and, thus, congenital VSMC differentiation defects in Tβ4-null mice may persist to adulthood and underlie medial degeneration. However, as previously reported, surviving embryos appeared to adequately compensate by normalising growth factor signalling to develop an overtly normal vasculature by embryonic day (E)14.5(7). We, therefore, examined aortas of postnatal day (P)7 male Tβ4^−/Y^ mice, compared with littermate Tβ4^+/Y^ controls (Figure 2A). Histological analysis of elastin integrity, aortic diameter, VSMC morphology and phenotypic marker expression confirmed that Tβ4^−/Y^ mice were indistinguishable from controls in the immediate postnatal period (Figure 2, A-C), suggesting a compensation during development. This raises the intriguing possibility that Tβ4 may be required throughout life to maintain vascular health and prevent defects in adulthood. To test this hypothesis, and to simultaneously determine if the postnatal requirement for Tβ4 is VSMC-autonomous, rather than paracrine, given the known roles in endothelial (6, 7) and immune cells(23, 24), we induced VSMC-specific loss of Tβ4 in postnatal mice. This was achieved by crossing a conditional (floxed) Tβ4 shRNA-expressing line (Hprt^Tβ4shRNA^; described in(8) (7)) with Myh11^CreERT2^(25) and administering three doses of tamoxifen (80mg/kg) to 3 week old mice, before examining their aortas at 12 weeks of age. Loss of *Tmsb4x* mRNA from aortic VSMCs was determined by RNA in situ hybridisation (RNAScope®; Figure 2D, quantified in Figure 2E). Compared with tamoxifen-dosed Myh11^CreERT2^; Hprt^+/+^ control mice, Myh11^CreERT2^; Hprt^Tβ4shRNA^ knockdown mice displayed a 1.8-fold aortic dilatation (mean, Figure 2, F-G), disruption of elastin lamellae (3.4-fold increase in number of breaks per section, Figure 2, H-J), only a modest degree of fibrosis (Figure 2K), and irregular VSMC morphology, with a shift towards expression of synthetic markers (Figure 2, L-N). The recapitulation of the global knockout phenotype suggests that reduction in Tβ4 levels through 8 weeks of postnatal life impairs vascular stability and defines a VSMC-autonomous, protective role in adult vessel homeostasis.

**Figure 2.**
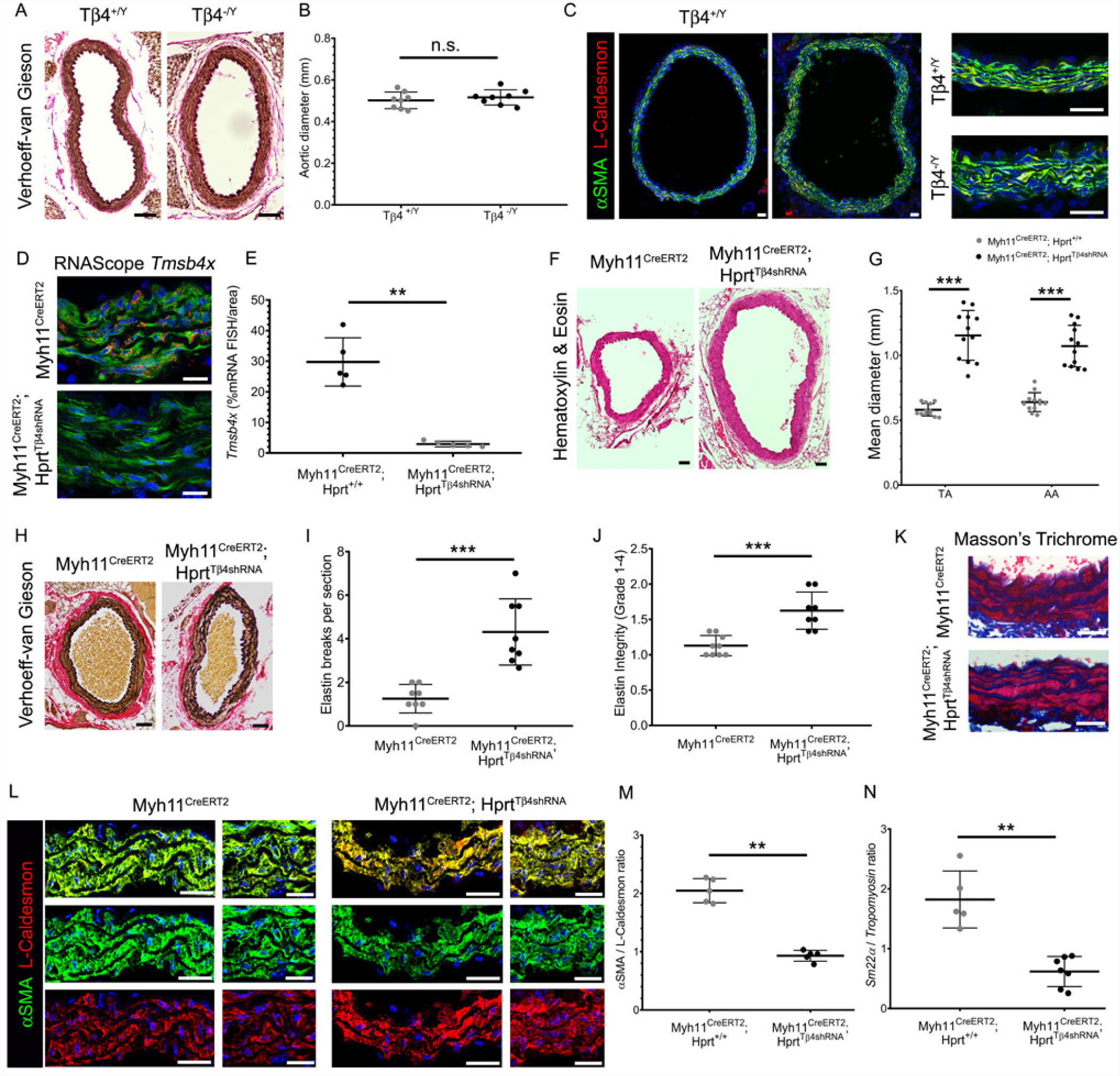
A postnatal, cell-autonomous requirement for Tβ4 in smooth muscle cells for maintenance of healthy aorta. Verhoeff-van Gieson staining (**A**) to visualise elastin deposition, structure and diameter (**B**) of aortas in postnatal day (P) 7 male mice. Immunofluorescence to assess smooth muscle phenotype(**C**). *Tmsb4x* was deleted from medial VSMCs of 3 week old Myh11^CreERT2^; Hprt^Tβ4shRNA^knockdown mice; RNAScope for *Tmsb4x* mRNA, compared with Myh11^CreERT2^; Hprt^+/+^control mice (**D**), quantified in **E**. Hematoxylin & Eosin staining (**F**) assessed aortic dilatation in 12-week old mice, quantified in **G**. Verhoeff-van Gieson staining (**H**) assessed elastin integrity, quantified both by number of breaks per section (**I**) and by an elastin damage score (**J**). Masson’s Trichrome staining (**K**) to examine collagen deposition. Ratio of contractile/synthetic VSMC markers in Myh11^CreERT2^; Hprt^Tβ4shRNA^, compared with Myh11^CreERT2^; Hprt^+/+^ aortas, both at the protein (**L**-**M**) and mRNA (qRT-PCR; **N**) level. Data are presented as mean ± SD, with each data point representing an individual animal. Significance was calculated using a Mann Witney non-parametric test (**B**, **E**, **M**, **N**) or two-tailed unpaired Student’s t-tests (**G**, **I**,**J**; with Holm-Sidak correction for multiple comparisons in **G**). n.s. = not significant; **p≤0.01; ***: p≤0.001. Scale bars: **A**, **F** and **H**: 100μm; **B**-**C, K**: 50μm; **D**: 20 μm; **L**: 50μm (low); 20μm (high).

### Tβ4 interacts with the vasculoprotective endocytic receptor, LRP1

To gain insight into the possible mechanisms by which Tβ4 maintains vascular health, we performed a yeast two hybrid screen to identify putative binding partners from an E11.5 murine embryonic library. A leading candidate was Low density lipoprotein receptor related protein 1 (LRP1), associated in human (12-14) and animal studies (15, 16) with protection against AAA and atherosclerosis. Two independent clones expressing regions of the intracellular domain of LRP1 were identified among the “prey” plasmids after stringent selection. Although we could not reproducibly co-immunoprecipitate Tβ4-LRP1 from murine aorta or MOVAS-1, a murine aortic VSMC line, we validated close association of the proteins (<40nm) by proximity ligation assay (PLA), consistent with the possibility of a weak or transient interaction, or an indirect association via a complex. Foci of Tβ4-LRP1 PLA signals were detected within medial VSMCs of murine aorta, at baseline (Figure 3A), as well as in disease (not shown). Moreover, conservation across species and clinical relevance was confirmed by detection of PLA signals in human VSMCs, both in aneurysmal aorta from AAA patients enrolled in the Oxford Abdominal Aortic Aneurysm Study (OxAAA study(26)); Figure 3B) and healthy vessels (matched omental artery samples from the same patients). Specificity for the PLA was ensured by lack of signal in -/Y Tβ4 null aortas (Figure 3A) and with omission of the LRP1 antibody (Figure 3B). While PLA signals were more abundant in omental arteries than AAA samples, inferences about disease relationship cannot be made, due to the small sample size and potentially compromised tissue integrity in the diseased vessels. Nevertheless, these observations collectively support a potential role for Tβ4 in regulating VSMC function via LRP1. We examined localisation of Tβ4-LRP1 foci more closely in primary murine aortic VSMCs. By immunofluorescence, LRP1 localised to punctate structures, consistent with its known incorporation into endocytic vesicles (Figure 3C). While Tβ4 was distributed throughout the cytoplasm, as expected, we observed strong puncta, suggesting that it may also localise to endosomal compartments. Indeed, Tβ4-LRP1 PLA signals partially overlapped with early endosomes, detected by expression of EEA1 (Figure 3D).

**Figure 3.**
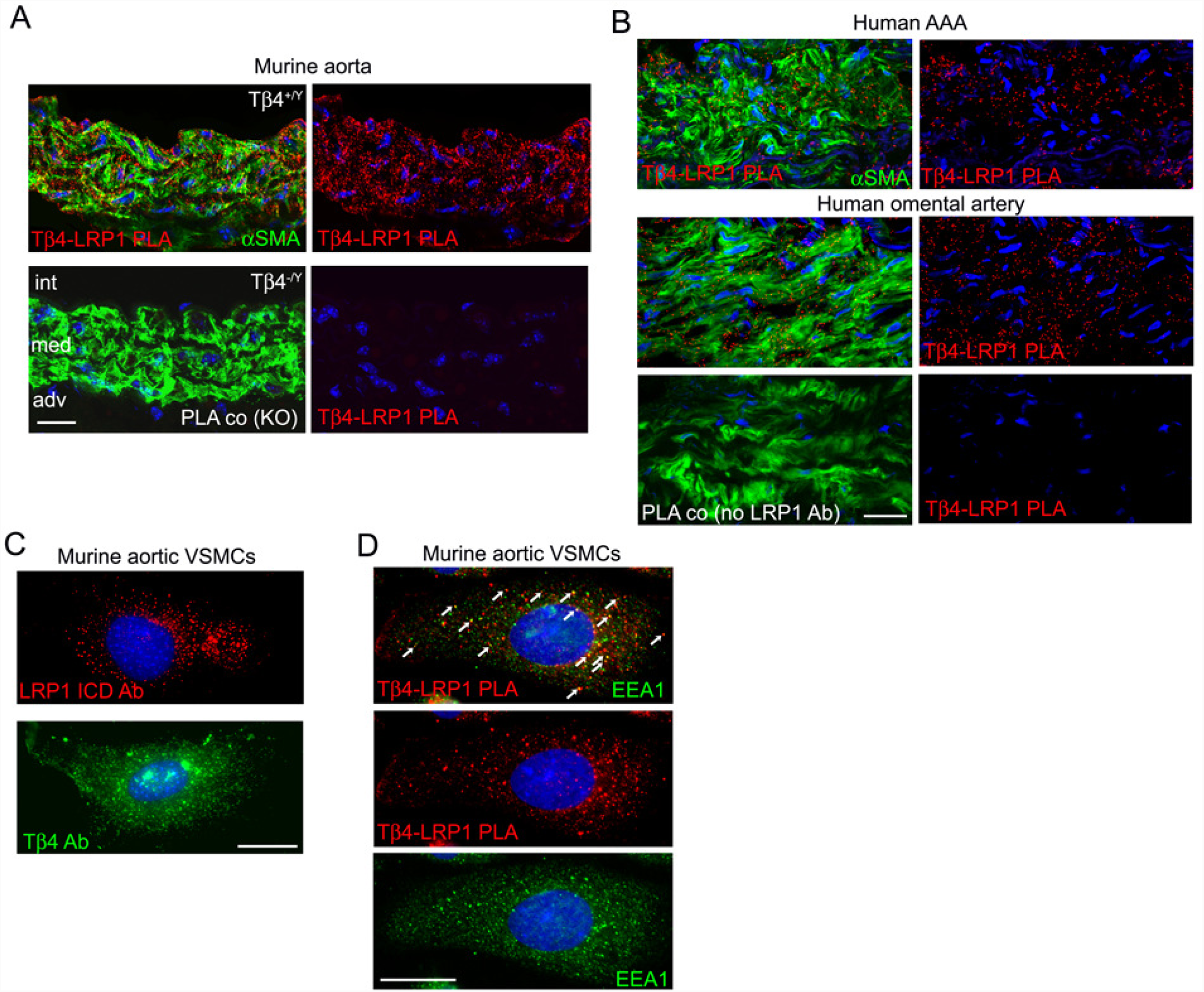
Tβ4 associates with Low density lipoprotein LDL receptor related protein 1 (LRP1) in endocytic vesicles of murine and human arterial smooth muscle cells. Proximity ligation assay (PLA) demonstrated close proximity (<40nm) and possible interaction of Tβ4 with LRP1 in murine aorta (**A**), human aorta, from AAA patients, and omental artery from the same patients (**B**). Representative images of n=3 (**A**-**B**). Immunofluorescence for LRP1 and Tβ4 reveals localization to punctate vesicular structures (**C**) in murine primary aortic VSMCs. PLA for Tβ4 and LRP1, with some signals localising to early endosomes, labelled with EEA1 (**D**). Scale bars: **A-B**: 20μm; **C-D**: 10μm. AAA: abdominal aortic aneurysm; PLA: proximity ligation assay; int: intima; med: media; adv; adventitia; ICD: intracellular domain (antigen).

### Loss of Tβ4 predisposes to vascular disease

Our assessment of the Tβ4^−/Y^ and Myh11^CreERT2^; Hprt^Tβ4shRNA^ vasculature at baseline reveals defects that phenocopy those previously reported in smooth muscle-specific LRP1^−/−^ mice(16, 17, 27) (aortic dilation, disrupted elastin layers and poorly differentiated VSMCs; Figures 1, 2 and comparison in Supplemental Figure 2). These observations not only support a common pathway, but also suggest that Tβ4KO mice may, like LRP1 nulls, be similarly predisposed to develop aortic aneurysm and atherosclerosis. We tested this supposition, firstly in a model of hypercholesterolaemia, by crossing onto the ApoE^−/−^ background(28) and feeding a ‘Western diet’ (WD; 21.4% fat; 0.2% cholesterol) for 12 weeks. Atherosclerotic plaques in the aortic sinus (Figure 4A-B) and descending thoracic aorta (Figure 4C) were significantly larger in Tβ4^−/Y^; ApoE^−/−^ mice than in Tβ4^+/Y^; ApoE^−/−^ littermate controls and were comparable in size to those of a smooth muscle-specific Lrp1 mutant (Myh11^Cre^; Lrp1^fl/fl^; ApoE^−/−^; tamoxifen-induced at 3 weeks of age). Increased plaque size did not correlate with an increase in weight gain or in cholesterol levels in Tβ4^−/Y^; ApoE^−/−^ mice, compared with Tβ4^+/Y^; ApoE^−/−^, although mice of both genotypes weighed more than Myh11^Cre^; Lrp1^fl/fl^; ApoE^−/−^, both at baseline and after high fat feeding (Supplemental Figure 3).

**Figure 4.**
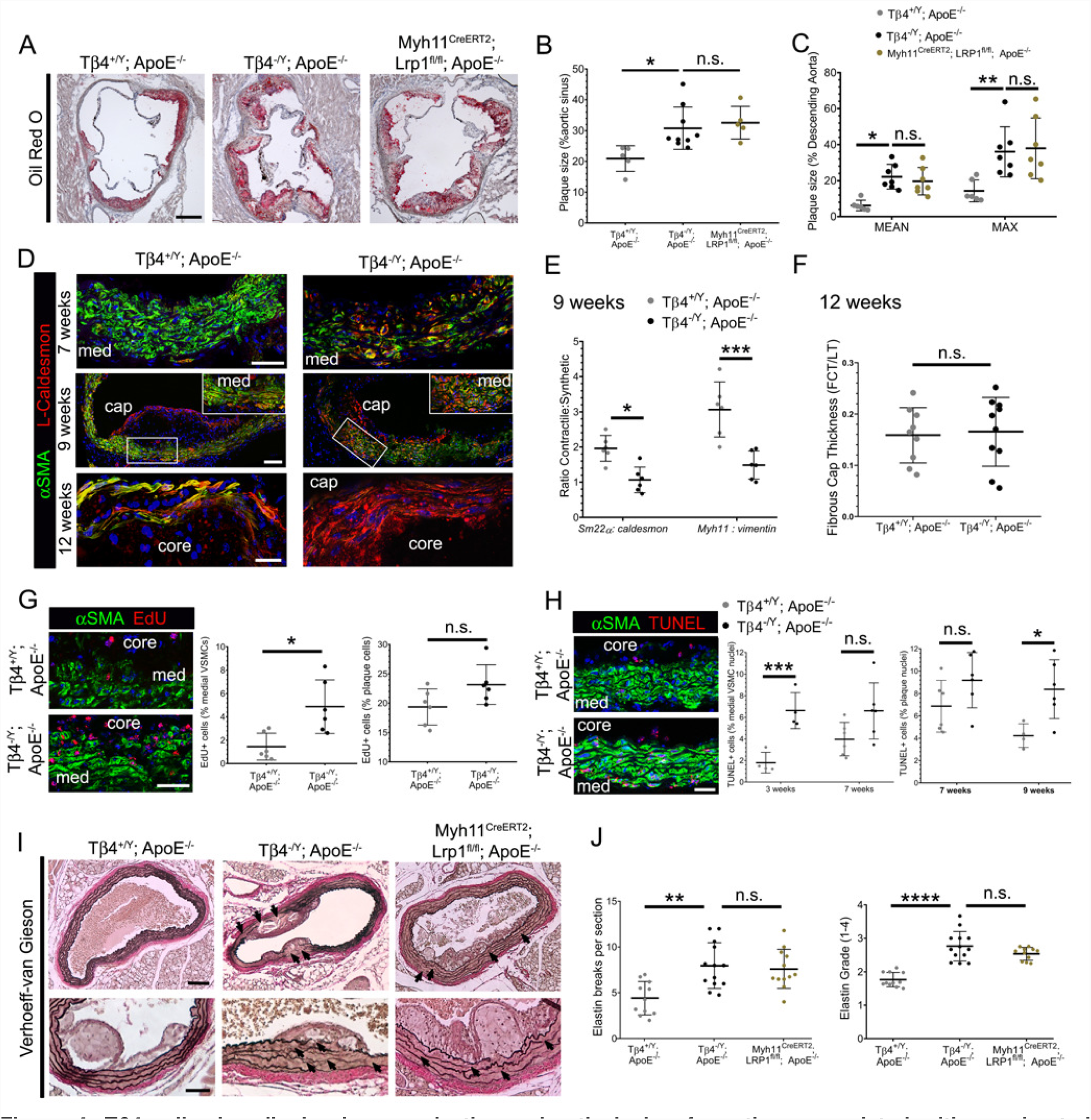
Tβ4-null mice display increased atherosclerotic lesion formation, associated with accelerated VSMC phenotypic switching. Oil red O staining of aortic sinus plaques (**A**), quantified in **B**. Descending aorta plaques quantified in **C**. Contractile-synthetic switching tracked over the course of lesion formation (**D**), quantified by qRT-PCR in **E**. Fibrous cap thickness quantified, relative to lesion thickness (**F**). Proliferation assessed by EdU injection (4 times over 10 days), prior to harvest at 7 weeks (**G**), with quantification of incorporation into medial VSCMs and plaque cells. Assessment of apoptotic medial VSMCs and plaque cells (TUNEL staining) over 3-9 weeks’ western diet feeding (**H**). Verhoeff-van Gieson staining of abdominal aorta to visualize elastin integrity underlying the plaque (**I**), with quantification of elastin breaks and degeneration (increased integrity score; **J**). Data are presented as mean ± SD, with each data point representing an individual animal. Significance was calculated using one-Way ANOVA with Tukey’s multiple comparison tests (**B**-**C**, **H, J**); Mann Witney non-parametric test (**E-G**). n.s. = not significant; *p ≤ 0.05; **p≤0.01; ***: p≤0.001; ****p≤ 0.0001. Comparisons are between Tβ4^+/Y^; ApoE^−/−^ and Tβ4^−/Y^; ApoE^−/−^mice and, where relevant, comparisons with Myh11^Cre^; Lrp1^fl/fl^; ApoE^−/−^aortas. Unless indicated, samples were harvested after 12 weeks’ Western diet. Scale bars: **A**:250μm; **D**: 50μm (upper); 100μm (middle); 20μm (lower). Boxed region in middle panel magnified in inset. **G-H**: 50μm; **I**: 200μm (upper); 100μm (lower). med: media; FCT: fibrous cap thickness; LT: lesion thickness.

We tracked VSMC phenotype over the course of WD feeding and, after 7 weeks, the medial layer VSMCs of Tβ4^−/Y^; ApoE^−/−^ mice displayed a pronounced alteration in morphology, with loss of their characteristic elongated shape and acquisition of a smaller, more rounded appearance(Figure 4D). This was accompanied by a shift in the ratio of synthetic (L-caldesmon and *vimentin* increased) over contractile markers (αSMA, *Sm22α* and *Myh11* decreased), determined by immunofluorescence and qRT-PCR (Figure 4D and E, respectively). Tβ4^+/Y^; ApoE^−/−^ VSMCs underwent a similar shift, only this occurred more gradually, becoming noticeable from 9 weeks’ WD, once plaques were already established (Figure 4D). We measured fibrous cap thickness, relative to lesion thickness, after 12 weeks and, whilst this was comparable between genotypes (Figure 4, D, F), the caps of Tβ4^−/Y^; ApoE^−/−^ plaques were composed of cells expressing exclusively synthetic markers, such as L-caldesmon (Figure 4D). In comparison, control Tβ4^+/Y^; ApoE^−/−^ cap cells retained higher levels of contractile markers, such as αSMA (Figure 4D). Consistent with dedifferentiation and acquisition of a synthetic phenotype, medial VSMCs were significantly more proliferative, as determined by increased EdU incorporation between weeks 5 and 7 of WD feeding (Figure 4G); in contrast, cells within the plaque showed no difference in proliferation between genotypes (Supplemental Figure 4A; quantified in Figure 4G). These cells, confirmed to be mostly macrophages/foam cells, based on CD68 expression (Supplemental Figure 4B), were present in similar number in Tβ4^−/Y^; ApoE^−/−^ and Tβ4^+/Y^; ApoE^−/−^ plaques, with plaque composition shown to be quantitatively indistinguishable, in terms of necrotic core and area occupied by macrophages (Supplemental Figure 4C). These data, along with comparable expression of pro-inflammatory cytokines, *Il1b* and *Tnfα* in descending aorta biopsies (Supplemental Figure 4D-E), suggest that altered inflammatory response are not primarily responsible for the altered medial layer phenotypes observed. Of note, a significantly higher proportion of Tβ4^−/Y^; ApoE^−/−^ medial VSMCs (3weeks’ WD) and plaque cells (9 weeks’ WD) were apoptotic, as revealed by TUNEL staining (Figure 4H), compared with Tβ4^+/Y^; ApoE^−/−^. Collectively, the preponderance of apoptotic and synthetic VSMCs, relative to the number of contractile VSMCs, suggests that, despite having thick fibrous caps, plaques in Tβ4^−/Y^; ApoE^−/−^ mice may be more unstable than those in Tβ4^+/Y^; ApoE^−/−^ mice. Finally, a striking degeneration of the elastin lamellae was apparent in Tβ4^−/Y^; ApoE^−/−^ aortas, with more elastin breaks per section than in Tβ4^+/Y^; ApoE^−/−^ aortas and a higher elastin damage score (Figure 4I-J). By these parameters, elastin degeneration in Tβ4^−/Y^; ApoE^−/−^ aortas was comparable with, and in some cases more severe than, Myh11^Cre^; Lrp1^fl/fl^; ApoE^−/−^ aortas (Figure 4I-J).

Whilst our study examined male mice in WD cohorts, the unanticipated observation of hindlimb paralysis, an indication of possible aortic thrombosis, aneurysm or rupture, prompted us to investigate a cohort of regular chow-fed 8 month old female Tβ4^−/−^; ApoE^−/−^ and age-matched Tβ4^+/+^; ApoE^−/−^ mice. Even without a cholesterol-rich diet, aortas of Tβ4^−/−^; ApoE^−/−^ mice were aneurysmal (>1.5 fold dilated), compared with Tβ4^+/+^; ApoE^−/−^, contained large atherosclerotic plaques and exhibited severe degeneration of the underlying elastin layers (Supplemental Figure 5). in view of the increasingly recognised sexual dimorphism in vascular disease, both in patients and in experimental studies(29), we compared a second cohort of female mice, this time with WD over an equivalent time course to that used in male cohorts, and determined a significant increase in aortic arch plaque size in Tβ4^−/−^; ApoE^−/−^, compared with Tβ4^+/+^; ApoE^−/−^ mice (Supplemental Figure 6). Interestingly, in female mice, as compared to male mice, increased plaque size did correlate with increased weight gain in Tβ4^−/−^; ApoE^−/−^ mice, compared with Tβ4^+/+^; ApoE^−/−^. Taken together, these studies suggest that atherogenesis is accelerated in Tβ4 null mice, regardless of gender, and is underpinned by an advanced VSMC contractile-synthetic phenotype shift, which closely recapitulates the phenotype associated with loss of LRP1.

We next determined whether loss of Tβ4 similarly predisposed to abdominal aneurysm, using the well-characterised model of Angiotensin II infusion (Ang II; 1mg/kg/day), via a subcutaneously implanted osmotic mini pump for up to 10 days. Analysis of whole mount aortas revealed an increased susceptibility of Tβ4^−/Y^ mice to aneurysm, compared with Tβ4^+/Y^ controls (Figure 5A). Phenotypes ranged from more pronounced ascending and descending aortic aneurysm (defined as >1.5-fold dilatation; mild) to abdominal aortic rupture, haematoma formation and death in <5 days (severe) (quantified in Figure 5B; examples of ruptures shown in Supplemental Figure 7). Although rare (<10%), dissections were detectable in Tβ4^−/Y^ as blood tracking into the adventitial matrix or between medial and adventitial layers; Supplemental Figure 7, death at day 8). Given the prominent role of inflammation in driving aneurysm progression with AngII treatment and numerous anti-inflammatory roles ascribed to Tβ4(23, 24, 30), we also investigated susceptibility in tamoxifen-dosed Myh11^CreERT2^; Hprt^Tβ4shRNA^ knockdown mice, alongside Myh11^CreERT2^; Hprt^+/+^ controls (Figure 5A) and Myh11^CreERT2^; Lrp1^fl/fl^ mice, in order to avoid disrupting Tβ4-LRP1 function in immune cells. Similar to global Tβ4 knockouts, VSMC-specific Tβ4 knockdown mice displayed an increased incidence of rupture (Supplemental Figure 7), as well as aortic aneurysms and a higher mortality rate over the 10 day time course (incidence rates quantified in Figure 5B and representative ruptures shown histologically in Supplemental Figure 7). Mean aortic diameter, measured on histological sections, was increased by 1.6-fold in Tβ4^−/Y^ mice and 1.8-fold in Myh11^CreERT2^; Hprt^Tβ4shRNA^ mice, compared with their respective AngII-treated controls. This compares with a 1.5-fold increased diameter in Myh11^Cre^; Lrp1^fl/fl^ aortas (Figure 5 C,D). Elastin integrity was severely breached in Tβ4^−/Y^, Myh11^CreERT2^; Hprt^Tβ4shRNA^ and Myh11^Cre^; Lrp1^fl/fl^ aortas, with more breaks per section and higher mean integrity scores than in Tβ4^+/Y^ and Myh^CreERT11^; Hprt^+/+^ controls (2.95, 2.85 and 2.82 versus 2.11 and 1.76, respectively; Figure 5 C,E). Beyond utilising a VSMC-specific targeting strategy, we sought to further exclude a major difference in inflammatory responses between genotypes by determining expression of pro-inflammatory cytokines *Tnfα, Il1b* and *Il6* by qRT-PCR in abdominal aortic biopsies and detected no significant difference between Tβ4^−/Y^ and Tβ4^+/Y^ (Supplemental Figure 8). Although VSMC degeneration is rapid in the AngII infusion model, we detected an accelerated VSMC dedifferentiation in Tβ4^−/Y^ (and Myh11^CreERT2^; Lrp1^fl/fl^) aortas, compared with respective controls (Figure 5F; shown after 5 days’ AngII). The prominent morphological alterations were accompanied by increased expression of synthetic maker, L-caldesmon (Figure 5F). After 10 days’ infusion, a further phenotype shift was observed, along with a greater VSMC loss, in Tβ4^−/Y^, Myh11^CreERT2^; Hprt^Tβ4shRNA^ and Myh11^Cre^; Lrp1^fl/fl^ aortas, compared with controls (Figure 5G). Collectively, these data demonstrate that loss of Tβ4 predisposes to aortic aneurysm, in the same way as loss of LRP1. Accelerated disease progression in mutants appeared not to relate to exacerbated inflammation, but rather to the more rapid VSMC phenotypic switch and degeneration of the elastin lamellae.

**Figure 5.**
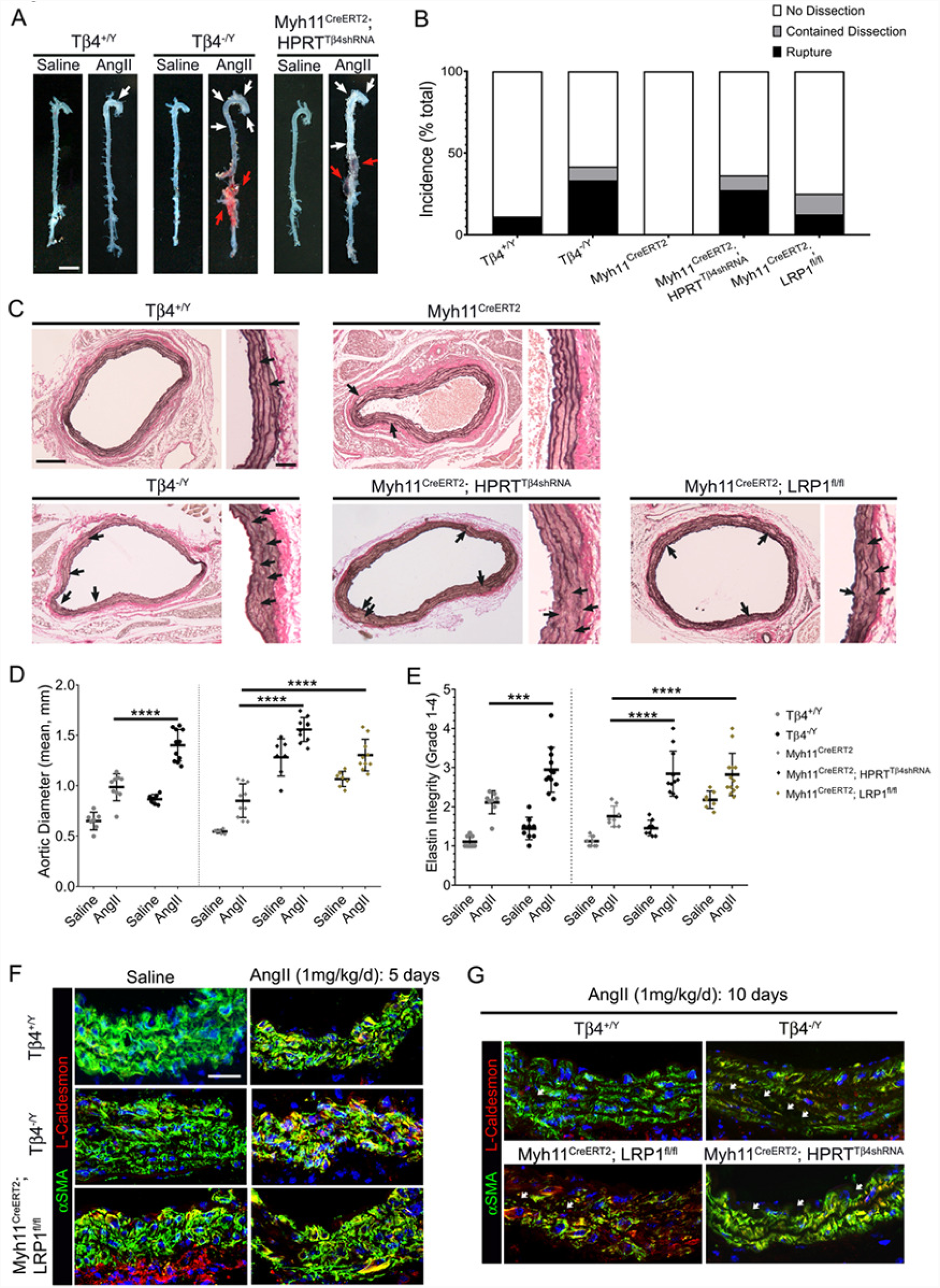
Mice lacking Tβ4 display more severe aneurysmal phenotypes, associated with accelerated VSMC phenotypic switching. Whole mount aortas from saline- or AngII-infused mice (**A**). White arrowheads indicate ascending and descending aortic aneurysm; red arrowheads indicate rupture. Incidence of dissection and rupture quantified per genotype in **B**. Verhoeff-van Gieson staining of abdominal aorta to visualize elastin integrity (**C**, breaks indicated with black arrowheads). Quantification of aortic diameter (**D**) and elastin grade (**E**). Immunofluorescence to assess medial layer morphology and VSMC phenotype at 5 (**F**) and 10 days’ (**G**) AngII infusion. Data are presented as mean ± SD, with each data point representing an individual animal. Significance was calculated using one-Way ANOVA with Tukey’s multiple comparison tests (**D-E**); ***: p≤0.001; ****p≤ 0.0001. Except in **F** (5 days), samples were harvested after 10 days’ AngII. Scale bars: **A**: 2mm; **C**: 500μm; inset 100μm.

### LRP1-controlled growth factor signalling is dysregulated in the absence of Tβ4

The VSMC-autonomous protective functions of LRP1 have been attributed, in part, to its role in regulating PDGFRβ endocytosis and signalling. We therefore sought further evidence for interaction of Tβ4 with the LRP1 pathway, by investigating readouts of the LRP1-PDGFRβ pathway in medial layer VSMCs early in the disease process, when injury-induced phenotypic switching is initiated. PDGF-BB binding to PDGFRβ stimulates autophosphorylation at multiple tyrosine residues; PDGFRβ tyrosine kinase activity requires LRP1-mediated endocytosis and, in turn, leads to phosphorylation of LRP1 on its intracellular domain (Tyr 4507)(31-33) to facilitate adaptor protein binding and activation of downstream pathways, including phosphatidylinositol 3-kinase (PI3K), Akt, p42/p44/MAP kinase (ERK1/2) and c-Jun N-terminal kinase (JNK)(34). After 3 weeks of WD, phospho-PDGFRβ (Tyr1021) and phospho-LRP1 (Tyr 4507) levels were elevated 2.4- and 1.8-fold, respectively, in VSMCs of Tβ4^−/Y^; ApoE^−/−^ aortas, compared with Tβ4^+/Y^; ApoE^−/−^ aortas (Figure 6A). Similarly, 1.8- and 2.8-fold increases in phospho-PDGFRβ (Tyr1021) and phospho-LRP1 (Tyr 4507), respectively, were observed in Tβ4^−/Y^ VSMCs, compared with Tβ4^+/Y^, after 5 days’ AngII infusion (Figure 6B).

**Figure 6.**
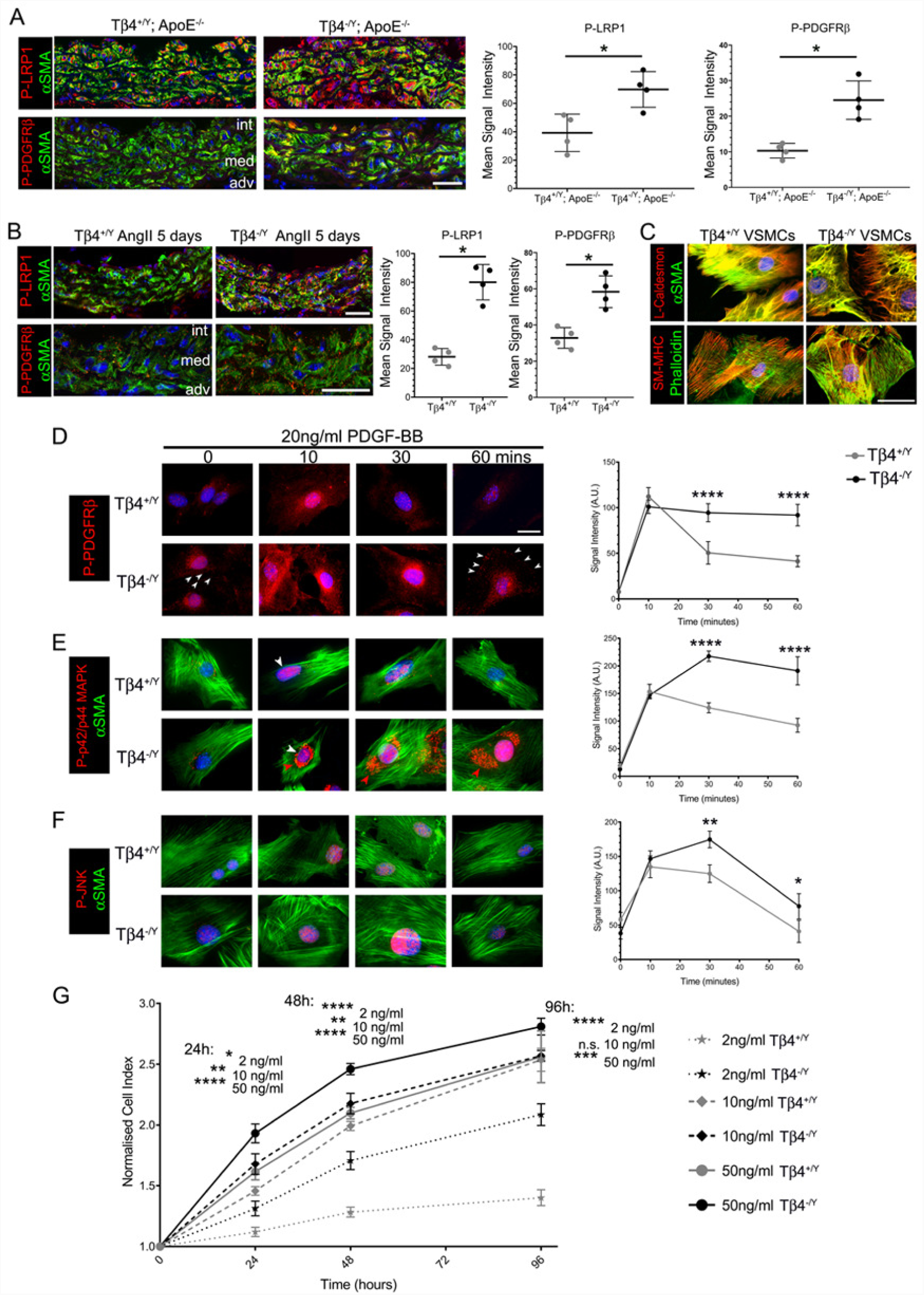
LRP1-PDGFRβ signalling is dysregulated in Tβ4-null VSCMs. Phospho-PDGFRβ (Tyr1021) and phospho-LRP1 (Tyr 4507) levels in atherosclerotic (**A**) and AAA (**B**) VSMCs. Isolated VSMCs from Tβ4^+/Y^ and Tβ4^−/Y^ aortas (**C**). Time course of PDGF-BB treatment and quantitative immunofluorescence of phosphorylated PDGFRβ (**D**) p42/p44 MAPK (ERK; **E**) and JNK (**F**). Proliferation curves of VSMCs treated with 2, 10 and 50ng/ml PDGF-BB (**G**). Data are presented as mean ± SD (**A**-**B**), with each data point representing an individual animal or mean ± SEM, n=3 experiments (**D**-**G**). Significance was calculated using a Mann Witney non-parametric test (**A**-**B**); two-Way ANOVA with Dunnett’s post-hoc tests (**D**-**G**). n.s. = not significant; *p ≤ 0.05; **p≤0.01; ***: p≤0.001;****p≤ 0.0001. Scale bars: **A**-**B**: 50μm; **C**: 50μm; **D** (applies **D**-**F**): 50μm.

Signalling responses of VSMCs were more closely interrogated *in vitro* in primary VSMC cultures established from the descending aortas of Tβ4^+/Y^ and Tβ4^−/Y^ mice. While some degree of VSMC dedifferentiation and contractile-synthetic phenotypic switching is inherent upon culture in high serum-containing medium, Tβ4^+/Y^ VSMCs typically exhibited a characteristic spindle-like morphology, whereas Tβ4^−/Y^ VSMCs were typically more rhomboid (Figure 6C). Although synthetic markers, such as L-caldesmon were expressed at comparable levels in the majority of VSMCs, expression of contractile VSMC markers, including αSMA and SM-MHC, was reduced in Tβ4^−/Y^ VSMCs. To compare signalling responses, serum-starved VSMCs were treated with 20ng/ml PDGF-BB over a 60 minute time course, prior to fixation, for analysis of pathway components by immunofluorescence. Phosphorylation (activation) of PDGFRβ, p42/p44 MAPK and JNK were strongly induced within 10 minutes of treatment (Figure 6, D-F). In Tβ4^+/Y^ VSMCs, phosphorylation of pathway components diminished by 30 minutes and returned to near-baseline by 60 minutes. This contrasted with Tβ4^−/Y^ VSMCs, in which phosphorylation remained high (PDGFRβ) or further increased (ERK, JNK) between 10 and 30 minutes and remained significantly higher (close to maximal Tβ4^+/Y^ levels) even at 60 minutes. These data reveal that PDGFRβ pathway activity is both enhanced in magnitude and more sustained in duration in Tβ4^−/Y^, compared with Tβ4^+/Y^ VSMCs, in response to the same dose of PDGF-BB. We, therefore, hypothesised that VSMCs lacking Tβ4 would be more sensitive to PDGF-BB-stimulated cellular responses, such as proliferation and migration. Indeed, over a range of tested PDGF-BB doses (2ng/ml – 50ng/ml), proliferation rate was significantly greater in Tβ4^−/Y^ than in Tβ4^+/Y^ VSMCs (Figure 6G). Of note, although only nuclear P-ERK signals were quantified above, a striking accumulation of perinuclear P-ERK was also apparent in Tβ4^−/Y^, but not Tβ4^+/Y^, VSMCs (Figure 6E); association of Tyrosine phosphorylated ERK with Golgi occurs during the G2/M phase of the cell cycle(35), further evidence of enhanced proliferation in Tβ4^−/Y^ VSMCs. By contrast, increased sensitivity to PDGF-BB did not correlate with enhanced migration, as assessed by scratch wound assay (Supplemental Figure 9); in fact, Tβ4^−/Y^ VSMCs migrated significantly more slowly than Tβ4^+/Y^ VSMCs. As this result was unexpected, we additionally compared migration rates in Myh11^Cre^; Lrp1^fl/fl^ VSMCs, which are similarly known to be hypersensitive to PDGF-BB, and found their migration to be likewise reduced compared with controls.

### Tβ4 modulates LRP1-PDGFRβ signalling via receptor-mediated endocytosis

As a proposed co-recepetor(31), endocytosis of LRP1 is necessary for PDGFRβ kinase activity(18). From endosomes, the LRP1-PDGFRβ complex may be recycled to the cell membrane or targeted for lysosomal degradation. Regulating receptor trafficking critically determines sensitivity to PDGF ligands and magnitude of downstream cellular responses. We, therefore, sought to investigate a role for Tβ4 in endocytosis of the LRP1-PDGFRβ complex, to determine if this may, at least in part, explain the effects of Tβ4 on PDGF-BB signalling described above. Initial studies were performed in MOVAS-1 cells, with siRNA-mediated knockdown of Tβ4 (reduced to 10.1 ± 7.9% at the mRNA level across all experiments; Supplemental Figure 10). Western blotting confirmed a similarly enhanced and more sustained activation of components of the PDGFRβ pathway in serum-starved, Tβ4 siRNA-treated MOVAS-1, with addition of 20ng/ml PDGF-BB (Figure 7A). Y1021-phosphorylated PDGFRβ and S473 phosphorylated Akt were significantly enhanced, although phosphorylated p42/p44 was unaffected in knocked down MOVAS-1 (Figure 7B). Total PDGFRβ levels declined steadily over the 60 minutes, consistent with lysosomal targeting and degradation and decline was notably more gradual in Tβ4 knockdown cells, albeit this was not significant (Figure 7B). In parallel, we performed surface biotinylation assays to measure levels of LRP1 and PDGFRβ at the cell surface over the same 60 minute time course of PDGF-BB treatment. Surface levels of LRP1 peaked after 5 minutes of treatment, with a comparable fold-change and rate of decline in knockdown and control cells up to 30 minutes. (Figure 7, C-D). However, whereas LRP1 levels in control cells declined further and remained significantly below baseline through to 60 minutes, consistent with the expected degradation(36), LRP1 levels in Tβ4 siRNA-treated cells recovered, almost to baseline, by 45 minutes (Figure 7C). Similar profiles were observed for PDGFRβ, except that the rate of decline in surface levels was more gradual in Tβ4 siRNA-treated cells, compared with controls. Levels fell between 5-45 minutes’ treatment; thereafter, a steep recovery of surface levels was observed in knockdown, but not in control, cells (Figure 7D). These data suggested that LRP1-PDGFRβ complexes are preferentially recycled to the cell surface in the absence of Tβ4 and that proportionally fewer receptors are targeted for lysosomal degradation.

**Figure 7.**
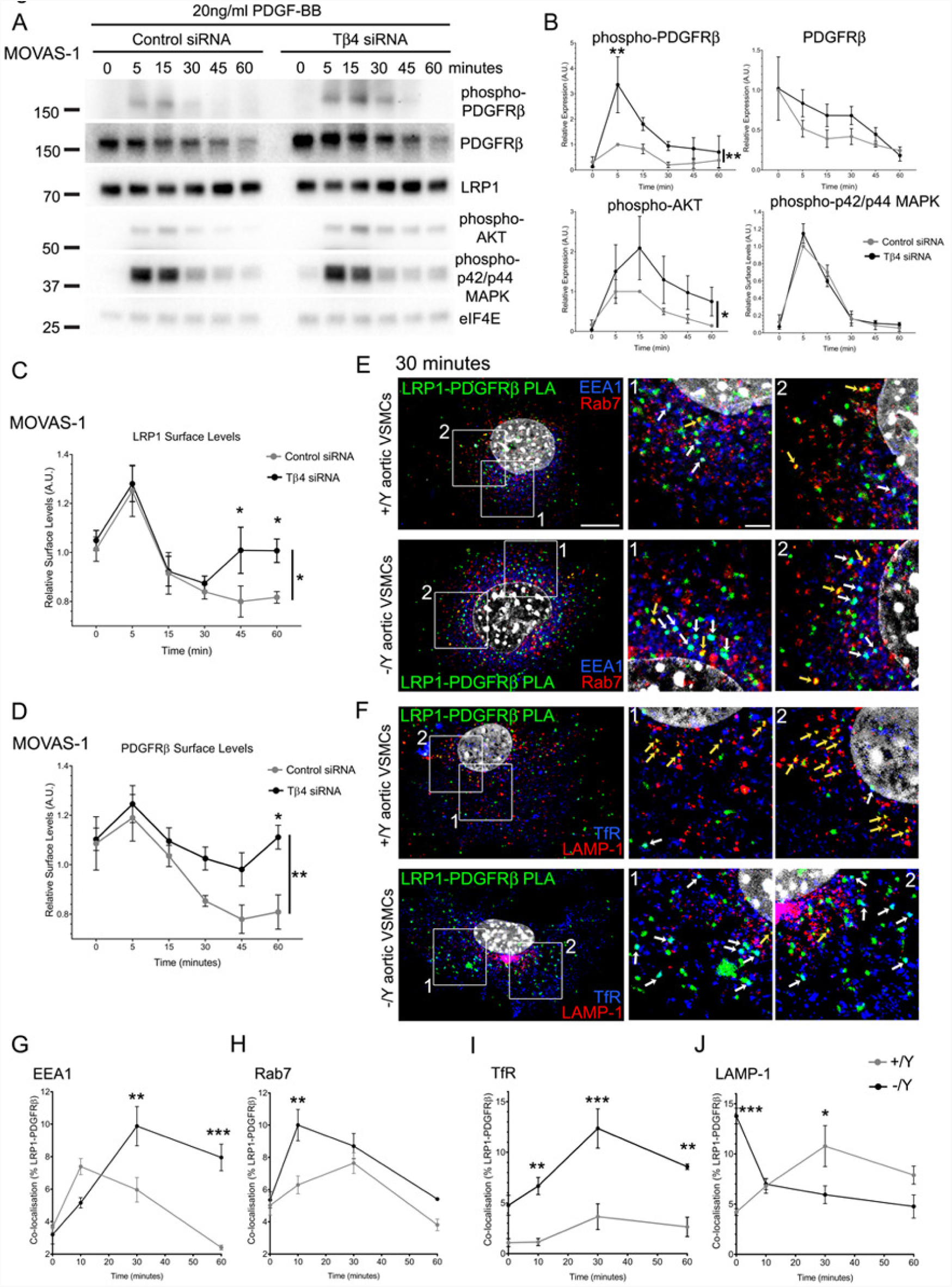
Loss of Tβ4 leads to dysregulation of LRP1-PDGFRβ receptor trafficking and enhanced sensitivity to PDGF-BB. Western blotting to assess PDGFRβ pathway activation in MOVAS-1 cells (**A**), quantified in **B**. Surface biotinylation assays measure levels of LRP1 (**C**) and PDGFRβ (**D**) at the cell surface. LRP1-PDGFRβ complexes, indicated by proximity ligation assay (PLA) in primary aortic VSMCs, colocalising with endocytic compartments over a 60 minute time course (**E**-**J**). Shown at 30 minutes, co-localising with early endosomes (EEA1; white arrows, **E**) and late endosomes (Rab7; yellow arrows, **E**) and with recycling endosomes (TfR; white arrows, **F**) and lysosomes (LAMP-1; yellow arrows, **F**). Quantification of co-localisation in **G**-**J.** Data are presented as mean ± SEM; n=4 experiments (**C**-**D**); n=3 experiments (**A-B**; **E**-**J**). Significance was calculated using two-way ANOVA with Sidak’s multiple comparison test (**B; C-D; G-J**).*p ≤ 0.05; **p≤0.01; ***: p≤0.001. Scale bars: **E:** 5μm (applies to all whole cell views in E-F); boxed areas 1 and 2 shown magnified to left, with scale 2μm (applies to all magnified views).

To test this notion further, we tracked the subcellular distribution of LRP1-PDGFRβ complexes (as PLA signals) through successive endocytic compartments over the 60 minute PDGF-BB time course, in primary murine aortic VSMCs from Tβ4^+/Y^ control and Tβ4^−/Y^ Tβ4 null mice (Figure 7E-J). Using antibodies that preferentially label early endosomes (EEA1), late endosomes (Rab7), recycling endosomes (Transferrin receptor; TfR) and lysosomes (LAMP-1), we quantified the proportion of LRP1-PDGFRβ colocalisation to each compartment using the ImageJ plug-in JACoP v2.0(37). In Tβ4^+/Y^ control cells, LRP1-PDGFRβ internalisation within EEA1+ early endosomes and Rab7+ late endosomes increased over the first 10 and 30 minutes’ treatment, respectively, before declining to below baseline levels (serum-starved cells; Figure 7, E, G-H). In Tβ4^−/Y^ cells, the proportions of LRP1-PDGFRβ in early endosomes increased further between 10 and 30 minutes and remained elevated in both early and late endosomes over the 60 minute time course, compared with control VSMCs (Figure 7, E, G-H). In Tβ4^+/Y^ control cells, only a small degree of LRP1-PDGFRβ recycling was observed (TfR+ recycling endosomes; Figure 7, F, I), and redistribution into LAMP-1+ lysosomes peaked after 30 minutes’ treatment (Figure 7, F, J). Recycling of LRP1-PDGFRβ in Tβ4^−/Y^ cells was significantly elevated at all time points, peaking at 30 minutes (Figure 7, F, I), and accompanied by a corresponding decline in levels of lysosomally targeted LRP1-PDGFRβ (Figure 7, F, J), albeit from an unexpectedly elevated baseline level in Tβ4^−/Y^ VSMCs, perhaps an adaptive response to suppress PDGF-BB hypersensitivity during conditions of serum starvation. Collectively, these findings suggest a requirement for Tβ4 in the attenuation of PDGFRβ signalling, following acute stimulation by PDGF-BB, via the targeting of LRP1-PDGFRβ complexes from late endosomes to lysosomes. A failure to down-modulate signalling may explain the advanced dedifferentiation (synthetic phenotype) and proliferation observed at baseline, which is further accelerated when PDGF-BB levels increase during disease.

## Discussion

Collectively, our results demonstrate a postnatal requirement for Tβ4 in VSMCs to maintain differentiated, contractile phenotype, both at baseline (homeostasis) and in the context of aortic diseases. Tβ4^−/Y^; ApoE-/-mice developed more unstable atherosclerotic plaques, in the aortic root and descending aorta. Similarly, global and VSMC-specific Tβ4 null mice displayed increased susceptibility to aortic aneurysm and a higher incidence of dissection, rupture and mortality. Accelerated disease progression was characterized by augmented contractile-synthetic VSMC switching and dysregulated PDGFRβ signaling, which may, at least in part, result from a failure to functionally regulate its endocytic co-receptor, LRP1. Consistent with this, the defects we describe in Tβ4 null mice closely phenocopy those reported with VSMC-specific loss of LRP1, both during development(38) and in disease(16, 17, 27). Of note, that the Myh11^CreERT2^; Lrp1^fl/fl^ mice described herein recapitulated the published LRP1 phenotype, even when loss of *Lrp1* was induced postnatally, confirms a maintenance role for LRP1, as with Tβ4, in vascular homeostasis, rather than persistence of developmental defects predisposing to disease. The previous studies used a constitutive sm22αCre(16, 27), active in embryonic VSMC progenitors and, as such, did not explicitly distinguish between these possibilities, although young mice were reported not to show aortic defects(17). Although we did not detect any overt exacerbation of inflammatory responses in global Tβ4^−/Y^ mice, it was important, due to the recognized roles for LRP1 and Tβ4 in endothelial cells(6, 39) and macrophages(40, 41), to distinguish a cell-autonomous VSMC role from potential paracrine contributions. However, it would be of interest to further investigate whether Tβ4 functionally regulates LRP1 in other cell types to influence vascular disease outcome.

From a clinical perspective, these findings are highly relevant. Genome-wide association studies have identified LRP1 variants as major risk loci for AAA(12), carotid artery (13) and coronary artery disease(14). Tβ4 is the most abundant transcript in healthy and AAA aorta(5), yet its roles in vascular protection and regulation of LRP1-mediated growth factor signalling had not been recognised. Our study provides a mechanism by which Tβ4 controls LRP1-mediated VSMC responses to protect against vascular disease. The significance of VSMC phenotypic switching in disease is still not fully resolved, as recently discussed in(42). While abnormal proliferation and migration of VSMCs drives early lesion formation and may contribute macrophage-like cells to the plaque, VSMCs are beneficial in advanced plaques, for preventing fibrous cap rupture. Understanding VSMC heterogeneity, and identifying the regulators of contractile-synthetic switching, may enable the fine tuning of VSMC phenotypes that are beneficial for repair. Interestingly, while loss of Tβ4 and LRP1 augmented PDGFRβ signaling and promoted proliferation, *in vitro* VSMC migration was, in our hands, inhibited. Although this finding is inconsistent with some studies demonstrating enhanced VSMC migration upon loss of LRP1(27, 43), others have demonstrated inhibition, confirming a requirement for LRP1 ((44, 45)), just as there is a clear requirement for Tβ4(46), in cell migration. Whether LRP1 promotes or inhibits migration of VSMCs appears to be dependent on extracellular matrix and ligand-binding cues (47), as discussed in(48). Moreover, there are likely distinct paracrine stimulatory roles for Tβ4-LRP1 as well as direct effects upon remodeling of the actin cytoskeleton(49) (50).

PDGF-BB, secreted from infiltrating macrophages during the initiating phases of aortic disease, potently drives VSMC phenotypic switching. The vasculoprotective effects of Tβ4 that we report appear to be mediated, at least in part, via LRP1-regulated growth factor signalling, specifically to control the sensitivity of VSMCs to PDGF-BB and potentially other ligands that are regulated by LRP1(51). Tβ4 was found to alter cellular responses to PDGF-BB by shifting the balance between receptor degradation and recycling. Further research is required to elucidate the precise mechanism(s) by which Tβ4 controls receptor trafficking and to determine whether this is limited to selected receptors, such as LRP1, or a generic regulation of endocytosis. *De novo* actin filament assembly is essential for endocytosis and, as an actin monomer-sequestering peptide, Tβ4 displays a biphasic effect: stimulating receptor-mediated endocytosis at low concentrations (<0.5μM) but inhibiting the process at high concentrations (10μM), consistent with actin depolymerization induced with high doses(52). More specifically, Tβ4-mediated, membrane-induced actin polymerization was shown to be required for fusion of late endosomes and phagosomes/lysosomes, but not for fusion of early endosomes, which can proceed without actin polymerization(53). Reduced lysosomal targeting of LRP1-PDGFRβ in our experiments is consistent with a defect in late endosome-lysosome fusion but seems an unlikely explanation for the increased partitioning to recycling endosomes. Recycling of signalling receptors occurs via a selective and carefully-regulated mechanism, which also requires actin polymerization, namely ‘Actin-Stabilized Sequence-dependent Recycling Tubule (ASSERT)’ scaffold formation(54), although a direct role for Tβ4 in this process has not been investigated. Thus, although there is support for a fundamental role for Tβ4 in endocytosis, given the remarkable similarity in developmental and disease phenotypes of Tβ4 and LRP1 mutants, a more selective role in regulation of LRP1-PDGFRβ by Tβ4 remains a possibility, supported by our identification of a physical interaction of LRP1-Tβ4 from a yeast two hybrid screen. While we failed to reproducibly validate the interaction by immunoprecipitation, GST pulldown and surface plasmon resonance (SPR), we cannot actually exclude a direct interaction, as there are technical complications with each of these approaches. The cytoplasmic domain of LRP1 interacts with multiple adaptor proteins and signalling proteins; while the 5kDa Tβ4 may participate, the complex may be too ‘congested’ to permit simultaneous binding of an anti-Tβ4 antibody, given the small size of the peptide. While pulldown and SPR methodologies obviate the need for antibody binding, they do not recreate the precise cellular context under which interaction potentially occurs, and which may be carefully regulated. Alternatively, an interaction may go undetected if it is transient and/or weak. A physical interaction is frequently inferred by proximity ligation assay, which requires proteins to be within 40nm and, in this regard, our data are supportive, although we cautiously conclude proximity, rather than a definitive protein-protein interaction. It is worth highlighting the functionally similar stabilin-2, an endocytic scavenger receptor which, like LRP1, internalizes coagulation factor VIII and LDL(55), amongst its many ligands, and contains NPxY motifs on its intracellular domain(56). Stabilin-2 was identified by yeast two hybrid(57) and validated by biolayer interferometry(58) to be a “weak and fuzzy”, yet specific, interactor of Tβ4, thus there is precedent for direct interaction with structurally and functionally similar receptors. Whether or not the proteins physically interact, we have validated an important functional interaction for Tβ4-controlled LRP1-PDGFRβ trafficking. Given the disease relevance of LRP1, not just in aortic disease but also in the pathogenesis of Alzheimer’s disease(59), further investigation into its internalization mechanism is warranted. Prevalence of aortic aneurysm is 5% amongst the elderly and treatment involves a high risk surgical procedure with no pharmacological therapeutic options. That Tβ4 can regulate endocytosis, even when administered exogenously(52), raises the possibility of developing novel strategies to maintain differentiated VSMC phenotype and treat aortic disease.

## Methods

### Animal models

Mice were housed and maintained in a controlled environment and all procedures involving the use and care of animals were performed in accordance with the Animals (Scientific Procedures) Act 1986 (Home Office, United Kingdom) and approved by the University of Oxford or University College London Animal Welfare and Ethical Review Boards. All mice used for the study were maintained on the C57BI6/J background for at least 10 generations. The global Tβ4 Knockout Tβ4(-/Y) strain, in which exon 2 of *Tmsb4x* locus is deleted, were a kind gift of Martin Turner, Babraham Institute and have been previously described(7). VSMC-specific Tβ4 knockdown mice were generated by crossing the previously described *Hprt1*-targeted, floxed Tβ4 shRNA line(7, 8) with tamoxifen-inducible Myh11^CreERT2^(25), to generate Myh11^CreERT2^; Hprt^Tβ4shRNA^. VSMC-specific Lrp1 knockouts were generated by crossing the previously described floxed Lrp1 line ((16); obtained from Jax) to homozygosity with Myh11^CreERT2^(25), to generate Myh11^CreERT2^; Lrp1^fl/fl^. These lines were crossed with ApoE^−/−^mice ((28); from Jax) to homozygosity. Conditional knockdown of Tβ4 or deletion of *Lrp1* were induced by oral gavage of 3 week old mice with 3 doses of tamoxifen (80mg/kg) on alternate days. Vascular injury studies were designed and executed in line with the recommendations of the American Heart Association Council on Arteriosclerosis, Thrombosis and Vascular Biology; and Council on Basic Cardiovascular Sciences(60).

### In vivo blood pressure measurement

One day after the MRI scan (exactly as previously described(61)), animals were anaesthetised with isoflurane (4%) and placed in supine position onto a heating plate with feedback control (Physitemp Instruments, NJ). Animals were kept at 37±1°C while oxygen and anaesthetics (1-2% isoflurane) were supplied via a nose cone (1 L/min). Body temperature and heart rate were recorded for the whole experiment using PowerLab with Chart 5 (ADInstruments, UK). A T-shaped middle-neck incision from mandible to the sternum was made. Blunt dissection was used to expose the left common carotid artery. A 5.0 Mersilk (Ethicon, NJ) suture was used to tie off the distal end of the artery while a micro vascular clip was used to occlude the proximal end of the exposed artery. A small incision near the distal end of the artery was made and a fluid filled tube (heparin; 100 U/mL diluted in 0.9% saline) with an inner diameter of 0.28 mm (Critchley Electrical Products Pty Ltd, Australia) was introduced into the artery and fixed in place with a suture. Following that, the micro vascular clip was removed and the arterial pressure was recorded using the MLT844 pressure transducer (ADInstruments, UK). After stabilisation of the signal for about 5-10 min an average of 1000 heart beats was used for analysis.

### Atherosclerosis model

Age-matched (8-12 week old) mice, on the Apoe^−/−^ background, were fed on Western diet (21.4% fat; 0.2% cholesterol; Special Diets Services, Essex, UK) for up to 12 weeks. Mice were weighed weekly and, at harvest, serum collected for cholesterol analysis (by Charles River Laboratories). EdU(5-ethynyl-2’-deoxyuridine), prepared in 0.9% saline, was administered at a dose of 150mg/kg by intraperitoneal injection (4 doses over the 10 days prior to harvest).

### Aneurysm model

8-12 week old adult mice were infused with 1mg/kg/day Angiotensin II (Sigma) by subcutaneous implantation of an osmotic mini pump (Alzet) for 10 days. Animals were regularly monitored and weighed post-surgery until harvest.

### Human tissue sampling

Subjects undergoing open abdominal aortic aortic aneurysm repair were prospectively recruited from the Oxford Abdominal Aortic Aneurysm (OxAAA) study. The study was approved by the Oxford regional ethics committee (Ethics Reference: 13/SC/0250). All subjects gave written informed consent prior to the study procedure. Baseline characteristics of each participant were recorded. During surgery to repair the AAA, a wedge of abdominal omentum containing a segment of omental artery was identified and biopsied en bloc. Isolation of omental artery was performed immediately in the operating theatre. The omental artery segment was cleared of perivascular tissue and snapped frozen. Prior to incision of the aortic aneurysm, a marker pen was used to denote the cross section of maximal dilatation according visual inspection. A longitudinal strip of the aneurysm wall along the incision was then excised. The aneurysm tissue was stripped off the perivascular tissue and mural thrombus. The tissue at the maximal dilatation was isolated, divided into smaller segments, and snap frozen for subsequent analysis.

### Histological Sample Preparation

For histological and immunohistochemical analyses, mouse aortic sinus, aortic arch and descending aorta were harvested, washed in phosphate-buffered saline (PBS; pH7.4) and fixed in 4% paraformaldehyde (PFA) at room temperature, either for two hours (cryosectioning) or overnight (paraffin embedding). For histological analysis, samples were dehydrated using a series of graded ethanol concentrations (50-100%, each for at least 2 hours), and cleared in butanol overnight. Samples were incubated at 60°C in 1:1 butanol: molten pastillated fibrowax for 30 minutes, then three times in 100% molten wax, each for 4 hours to overnight, before embedding and sectioning (10μm transverse sections). Paraffin sections were stained with Oil red O solution, Hematoxylin and Eosin, Elastic stain kit (Verhoeff van Gieson) and Masson’s Trichrome kit (all from *Sigma-Aldrich*), each according to the manufacturer’s protocol. After imaging, morphology was assessed on sections which had been anonymised and blinded for genotype. Elastin breaks per section were counted manually. Plaque area and aortic diameter were measured using ImageJ. Elastin breaks/score, aortic diameter, TA/AA plaque size are all expressed as the mean of 6 sections per aorta) For immunohistochemical analyses, fixed tissues were rinsed in PBS and equilibrated in 30% sucrose (in PBS) overnight at 4°C. Samples were placed in 50:50 (30% sucrose: Tissue-Tek OCT), then embedded in OCT and chilled at - 80°C.

### Immunofluorescence Staining on Cryosections

Frozen sections of descending aorta or aortic sinus (base of the heart) were cut at a thickness of 10 μm, air-dried for 5 minutes and rinsed in PBS. Sections were permeabilized in 0.5% Triton-X100/PBS for 10 minutes, rinsed in PBS and blocked for 1-2 hours (1% BSA, 10% goat serum or donkey serum; 0.1% Triton-X100 in PBS). Sections were incubated with primary antibodies (diluted as below) at 4°C overnight. Sections were washed 5-6 times in 0.1% Triton-X100 in PBS (PBST) then incubated in secondary antibody for 1 hour at room temperature. Sections were washed 3 times in PBST and, incubated with 300nmol/uL 4’,6-diamidino-2-phenylindole (DAPI) in PBS for 5 minutes and rinsed a further twice in PBS. Slides were mounted with 50% glycerol:PBS and imaging was conducted on an Olympus FV1000 confocal microscope or a Leica DM6000 fluorescence microscope. Apoptosis was assessed using the Click-iT TUNEL Alexa Fluor 546 Imaging Assay kit and EdU incorporation by Click-iT EdU Alexa Fluor 546 Imaging Assay kit (both from ThermoFisher), according to the manufacturers’ instructions.

### Cell Culture

Primary smooth muscle cells were isolated by enzymatic digestion of murine aortas, as previously described(62). After the first week, fetal bovine serum (FBS) was gradually reduced from 20% to 10% and cells maintained thereafter in DMEM (Gibco) supplemented with 10% FBS, 100 U/ml Penicillin-Streptomycin (Gibco), in a humidified incubator at 37°C and 5% CO_2_. An immortalised mouse vascular aortic smooth muscle cell line (MOVAS-1) was purchased from ATCC and cultured in DMEM (ATCC) supplemented with 10% FBS, 100 U/ml Penicillin-Streptomycin (Gibco) and 0.2 mg/ml G −418 (Sigma) at 37°C and 5% CO_2_.

### siRNA treatment

MOVAS-1 cells were transfected with 15 nM siRNA against Thymosin β4 (s201860, Thermo Scientific) or control siRNA (Silencer Select negative control, 4390844, Thermo Scientific) using Lipofectamine RNAiMAX (Thermo Scientific) for 24-36 hours. In parallel, siRNA-treated cells were starved in medium without FBS prior to stimulation with 20 ng/ml PDGF-BB for fixed periods of time and processed for Western blot, ELISA or Duolink, as described.

### Antibodies for Immunofluorescence

Cy3 conjugated Mouse anti-αSM actin (1:200; Sigma; C6198); FITC conjugated Mouse anti-αSM actin (1:200; Sigma; F3777); Rabbit anti-Vimentin (1:200; Abcam; ab45939); Rat anti-CD68 (1:100; BioRad; MCA1957); Rabbit anti-SM-MHC (1:1000; Abcam; ab125884); Rat anti-Ki67 (1:100; eBioscience; 14-5698); Rabbit anti-Caldesmon (1:250 Abcam; ab32330); Rabbit anti-Thymosin β4 (1:100; Immunodiagnostik; A9520); Mouse anti-LRP1 ICD hybridoma monocolonal antibody (1:25; ATCC CRL-1937); Rabbit anti-phospho-p44/42 MAPK (Erk1/2) (Thr202/Tyr204) (1:200; Cell Signalling Technology, 4370); Rabbit anti-phospho-LRP1 (Tyr4507) (1:100; Santa Cruz Biotechnology, sc-33049); Rabbit anti-phospho-PDGFRβ (Tyr1021) (1:100; Abcam, ab134048); Rabbit anti-phospho JNK1 + JNK2 + JNK3 (T183+T183+T221) (1:100; Abcam, ab124956); Armenian Hamster anti-PECAM1 (1:100; Abcam, ab119341); Cy3-conjugated Rabbit anti-LAMP1 (1:25; Abcam, ab67283), Rat anti-Transferrin receptor (1:25; Abcam, ab22391), AlexaFluor 647-conjugated Rabbit anti-EEA1 (1:50; Abcam, ab2900) and AlexaFluor 647-conjugated Rabbit anti-RAB7 (1:100; Abcam,ab198337). AlexaFluor-conjugated secondary antibodies raised against Rat, Rabbit, Mouse or Armenian Hamster IgG (Invitrogen) were used at 1:200.

### Immunoblotting

SiRNA-treated and PDGFβ-stimulated cells were lysed in RIPA buffer (50 mM Tris-HCl pH7.5, 100 mM NaCl, 1% NP-40, 0.1% SDS, 0.5% sodium deoxycholate, PhosSTOP and cOmplete protease inhibitor cocktail (Roche)). Western blots were performed using a standard protocol. Proteins were transferred by semi-dry transfer onto PVDF membrane. After blocking, membranes were probed with primary antibodies overnight. A modified protocol was used for Thymosin β4 Western blotting(63). Proteins were separated on Tris-Tricine gels and cross-linked with 10% glutaraldehyde, before wet overnight transfer onto nitrocellulose membrane. Membranes were crosslinked with UV light (254 nm, 10 min) before blocking and overnight antibody probing. Antibodies used were Rabbit anti-phospho-PDGFRβ (Tyr1021) (1:500; Abcam, ab62437), Rabbit anti-PDGFRβ (1:1000; Abcam, ab32570), Rabbit anti-LRP1 (1:10,000; Abcam, ab92544), Rabbit anti-phospho-Akt (S473) (1:500; Cell Signalling Technology, 4058), Rabbit anti-phospho-ERK1/2 (Thr202/Tyr204) (1:1000; Cell Signalling Technology, 9101), Rabbit anti-eIF4E (1:1000; Cell Signalling Technology, 2067), Rabbit anti-GAPDH (1:1000; Proteintech, HRP-60004) and Sheep anti-Thymosin β4 (1:100; R&D, AF6796). HRP-conjugated secondary antibodies were from GE Healthcare (rabbit, NA934) and R&D (sheep, HAF016).

### Duolink® Proximity Ligation Assay

Aortic sections, primary VSMCs or SiRNA-treated and PDGF-BB-stimulated MOVAS-1 cells were fixed with 4% PFA and permeabilised with 0.1% Tween-20. To assess proximity, the following antibody combinations were used: Mouse anti-LRP1 (1:100; Abcam, ab28320) and either Rabbit anti-PDGFRβ (1:100; Abcam, ab32570) or Rabbit anti-Thymosin β4 (1:100; Immundiagnostik, aa1-14). Duolink***®*** In Situ PLA Rabbit Plus and Mouse Minus probes were used in combination with Detection Reagent Green (Sigma) following manufacture’s protocol. Following Duolink***®*** protocol, cells were co-stained for smooth muscle markers or endocytic compartments using the antibodies described above. Colocalisation was determined, using the ImageJ plug-in JACOP 2.0(37).

### Surface biotinylation ELISA

SiRNA-treated and PDGF-BB-stimulated cells were washed twice with cold PBS and surface proteins were biotinylated with 0.5 mg/ml Sulfo-NHS-SS-Biotin (Thermo Scientific) for 30 min on ice. The reaction was quenched with 15 mM glycine (2x 5 min) and cells were lysed in GST lysis buffer (10% glycerol, 50 mM Tris-HCl pH7.5, 200 mM NaCl, 1% NP-40, 2 mM MgCl_2_, PhosSTOP and cOmplete protease inhibitor cocktail (Roche)). Nunc-Immuno™ MicroWell™ 96 well plates were coated with antibody in 0.05 M Na_2_CO_3_ pH9.6 (anti-LRP1 (1:1000, Abcam, ab92544) or anti-PDGFRβ (1:500, Abcam, ab32570)) overnight at 4°C. Plates were washed and blocked for 1 hour in 1% Hammarsten grade casein (Alfa Aesar) with 0.1% Tween-20 in PBS. Lysates were quantified by BCA assay, diluted in blocking buffer, equal amounts added to the plate and incubated overnight. Plates were washed, incubated with streptavidin-HRP (1:1000, Abcam, ab7403) for 1 hour and developed with TMB substrate (BD Biosciences). The reaction was stopped with 1M sulphuric acid and absorbance read at 450 nm. ***RNA isolation and gene expression profile by quantitative real time (qRT-PCR)***. Total RNA was isolated from the aortic arch or descending aorta of adult mice using the RNeasy Fibrous Tissue Mini kit (Qiagen), according to the manufacturer’s instructions. cDNA was prepared using RT kit (Applied Biosystems). qRT-PCR was performed on a ViiA 7 (Life technologies) using fast SYBR Green (Applied Biosystems). Data were normalised to β-actin expression (endogenous control, after selection of optimal control, in terms of stable expression, from a panel of 6 genes). Fold-changes were determined by the 2-^ΔΔ^CT method(64).

### Primers for qRT-PCR (5’-3’)

*b-actin* F: GGCTGTATTCCCCTCCATCG; R: CCAGTTGGTAACAATGCCATGT; *Acta2* F: GTCCCAGACATCAGGGAGTAA; R: TCGGATACTTCAGCGTCAGGA; *sm22α* F: CAACAAGGGTCCATCCTACGG; R: ATCTGGGCGGGCCTACATCA; *Myh11* F: AAGCTGCGGCTAGAGGTCA; R: CCCTCCCTTTGATGGCTGAG; *Caldesmon* F: GCTCCCAAGCCTTCTGACTT; R: CCTTAGTGGGGGAAGTGACC; *Tropomyosin IV* F: CGCAAGTATGAGGAGGTTGC; R: AGTTTCAGATACCTCCGCCCT; *Vimentin* F: TCCAGCAGCTTCCTGTAGGT; R: CCCTCACCTGTGAAGTGGAT; *Il1b* F: CAACCAACAAGTGATATTCTCCATG; R: GATCCACACTCTCCAGCTGCA; tnf*α* F: CACGTCGTAGCAAACCACCAAGTGGA; R: TGGGAGTAGACAAGGTACAACCC

### RNAscope® single molecule RNA in situ hybridisation

The RNAscope® Multiplex Fluorescent v2 Assay (AcDBio) was used to visualize and quantify single molecule mRNA targets per cell on cryosections. Fixed aortic sections were pre-treated with RNAscope Pre-treatment kit to unmask target RNA and permeabilise. Protocol was performed according to the manufacturer’s instructions, using the RNAscope® Probe - Mm-Tmsb4x-C2 472851-C2, followed by sequential amplification of signal using Opal detection reagents (Invitrogen). Sections were imaged using an Olympus FV1000 confocal microscope.

### Smooth muscle proliferation assays

After trypsinization, 1000 cells/well were plated in DMEM; 10% FBS into 96-well plates. After overnight serum starvation, medium was replaced with serum-free DMEM containing PDGF-BB (2-50ng/ml). At the harvest time point, plates were washed frozen and processed for DNA-based assay, as described(65), and background subtracted using a plate which had been incubated with medium but no cells.

### Smooth muscle migration assays

Primary aortic VSMCs were cultured as a confluent monolayer in 12-well plates. After overnight serum starvation, scratch wounds were introduced with a P200 tip. After washing, medium was replaced with serum-free DMEM containing PDGF-BB (2-50ng/ml) and 100 μM hydroxyurea (Sigma). Wells were imaged at t0, and every 24h thereafter, using a Zeiss Axio Imager microscope. Wounds were measured and expressed as % relative to t0.

### Statistics

Randomisation of animals to treatment or genotype groups was introduced at the time of mini pump implantation (aneurysm); introduction of western diet (atherosclerosis) or harvest (baseline). Thereafter, tissues were processed and analysed by an independent observer whilst blinded to treatment or genotype. Statistical analyses were performed with GraphPad Prism Software. For the quantitative comparison of two groups, two-tailed unpaired Student’s t-test was used to determine any significant differences, after assessing the requirements for a t-test using a Shapiro-Wilk test for normality and an F-Test to compare variances. Alternatively, a Mann-Whitney non-parametric test was used. For comparison of three groups or more, a one-way ANOVA with Tukey’s post-hoc test was used. For analyses involving two independent variables, a two-way ANOVA with Bonferroni, Holm-Sidak or Dunnett’s post-hoc test was used. Significance is indicated in the figures, as follows: *: p≤0.05; **: p≤0.01; ***: p≤0.001; ****: p≤0.0001.

### Study approval

The Oxford Abdominal Aortic Aneurysm (OxAAA) study was approved by the Oxford regional ethics committee (Ethics Reference: 13/SC/0250). All procedures involving the use and care of animals were performed in accordance with the Animals (Scientific Procedures) Act 1986 (Home Office, United Kingdom) and approved by the University of Oxford or University College London Animal Welfare and Ethical Review Boards.

## Author contributions

SM, SB and NS carried out most experiments and data analysis, with additional data contributed by ANR, KND, RD and GN. JP and ANR performed surgical procedures. RL and AH designed and conducted the OxAAA study, which enabled the human tissue analyses. KMC provided valuable intellectual input. NS established the hypotheses, supervised the study and wrote the manuscript.

## Acknowledgements

The study was funded primarily by the British Heart Foundation Ian Fleming Senior Basic Science Research Fellowship (FS/13/4/30045), awarded to NS, and also by a studentship from the Oxford Medical Research Council Doctoral Training Partnership to SM.

