## Supplemental Figures 1-10 for "Thymosin β4 mediates vascular protection via regulation of Low Density Lipoprotein Related Protein 1 (LRP1)"

### Online Supplemental Data

Supplemental Figures 1-10

Elastin Integrity: Scoring Grades 1-4

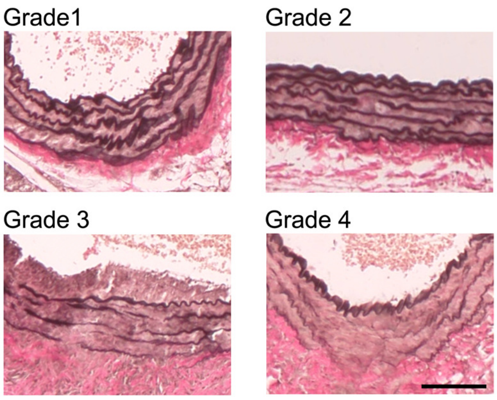


**Supplemental Figure 1. Visual scoring system used to assess severity of elastin degradation.** Grades 1-4 relate to increasing severity (1=normal; 2=occasional breaks; 3= frequent breaks, multiple lamellae; 4=severe breakdown). Aortas were scored, whilst blinded to genotype. 6-8 sections per aorta were assessed and mean score plotted for each animal. Scale bar (applies to all panels): 100µm.

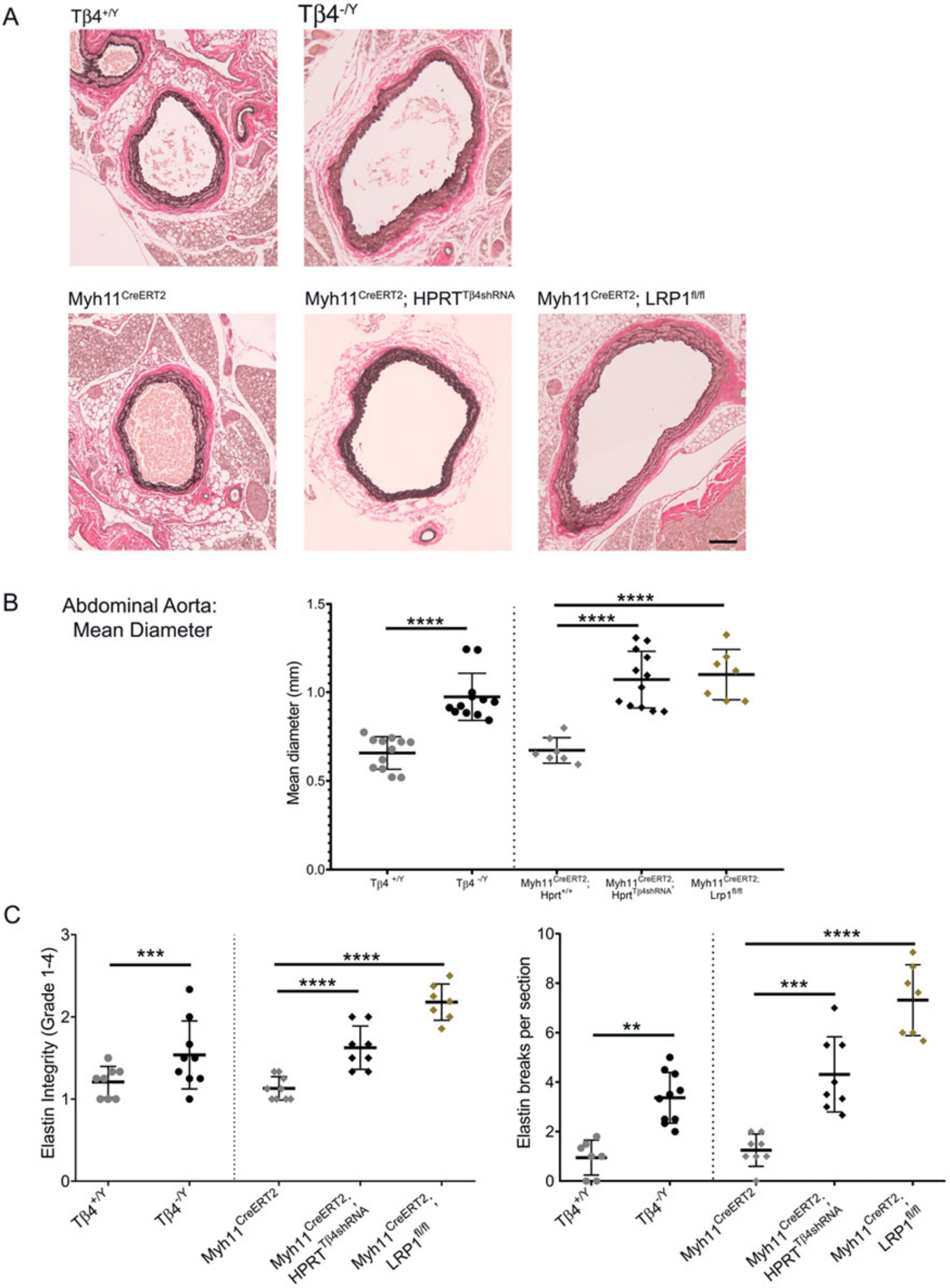


**Supplemental Figure 2. Global and VSMC-specific loss of Tβ4 phenocopies VSMC-specific *Lrp1* mutants.** Baseline phenotypes of $T\beta4^{-/Y}$, compared with $T\beta4^{+/Y}$, and $Myh11^{CreERT2}$; $Hprt^{T\beta4shRNA}$ and $Myh11^{CreERT2}$; $Lrp1^{fl/fl}$, compared with $Myh11^{CreERT2}$, revealed by Verhoeff-van Gieson staining (**A**), quantification of aortic diameter (**B**) assessed elastin integrity, quantified both by number of breaks per section and by an elastin damage score (**C**). Significance was calculated using one-Way ANOVA with Tukey's multiple comparison tests (**B**-C). **p≤0.01; ***: p≤0.001; ****p≤ 0.0001. Scale bar: **A**: 100µm.

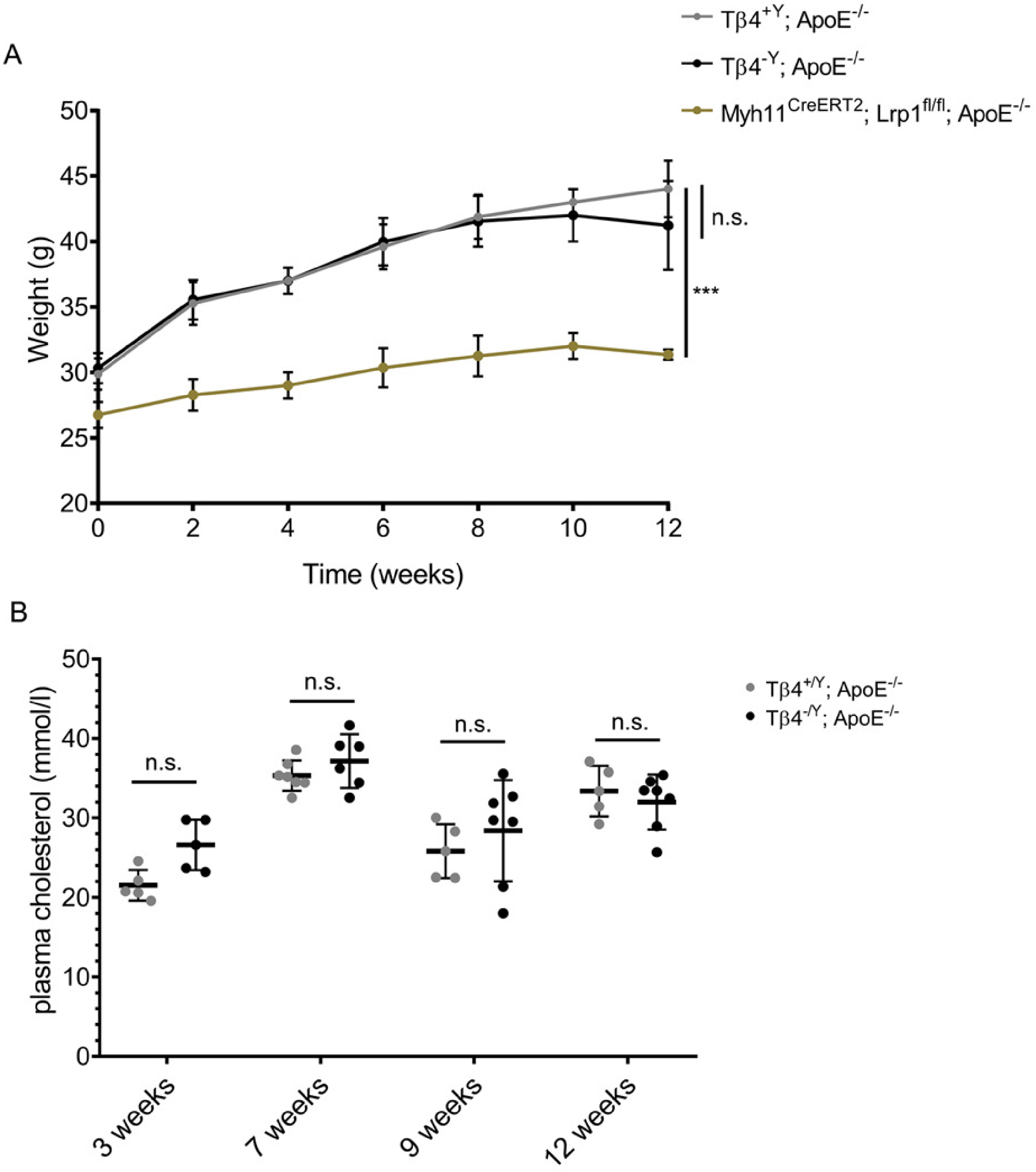


**Supplemental Figure 3. Tβ4$^{+/Y}$; ApoE$^{-/-}$ and Tβ4$^{-/Y}$; ApoE$^{-/-}$ do not show differences in weight gain or cholesterol levels after western diet.** Comparison of weight gain (**A**) and plasma cholesterol levels (**B**) in Tβ4$^{+/Y}$; ApoE$^{-/-}$ and Tβ4$^{-/Y}$; ApoE$^{-/-}$ mice over the time course of western diet feeding regime. Data are presented as mean ± SD, with n=9-11 in **A**. Each data point in **B** represents an individual animal. Significance was calculated using two-way ANOVA with Bonferroni correction for multiple comparisons (**A**) and one-Way ANOVA with Tukey's multiple comparison tests (**B**). n.s. = not significant; ***: p≤0.001

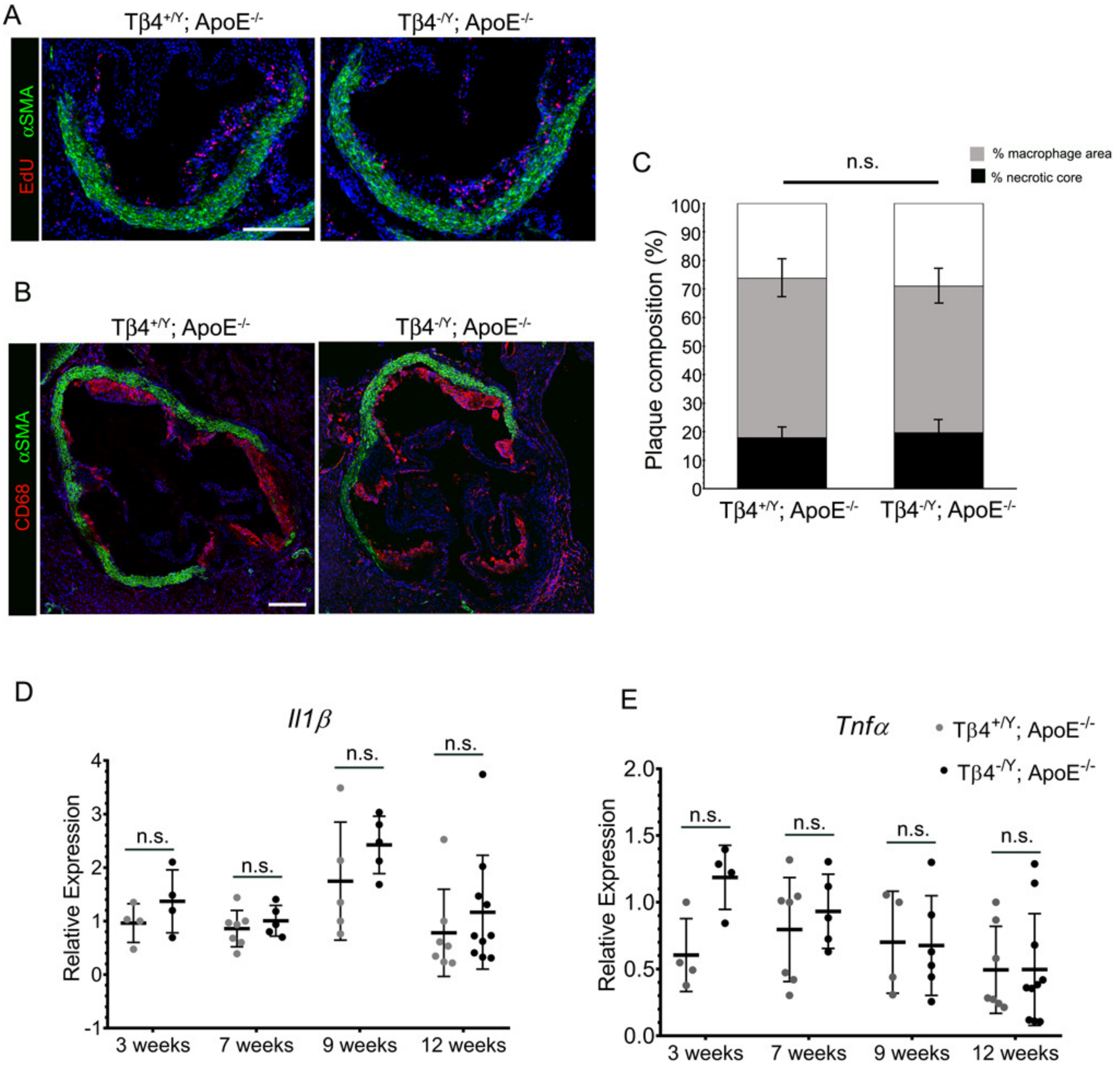


**Supplemental Figure 4. Inflammatory responses in atherosclerosis are not significantly altered with loss of Tβ4.** Low power image to demonstrate that most EdU+ cells in the aortic sinus at 7 weeks (EdU doses between 5-7 weeks) were present within (**A**). Immunofluorescence demonstrates CD68+ macrophage/foam cells in the plaque (**B**). Plaque composition, in terms of % occupied by macrophages and % necrotic core (determined from histological staining; **C**). qRT-PCR of pro-inflammatory cytokine expression: *Il1β* (**D**) and *Tnfα* (**E**). Data are presented as mean ± SD, with each data point representing an individual animal. Significance was calculated using Mann Witney non-parametric test (**C**) one-Way ANOVA with Tukey's multiple comparison tests (**D**-**E**). n.s. = not significant. Scale bar: **A-B**: 250µm.

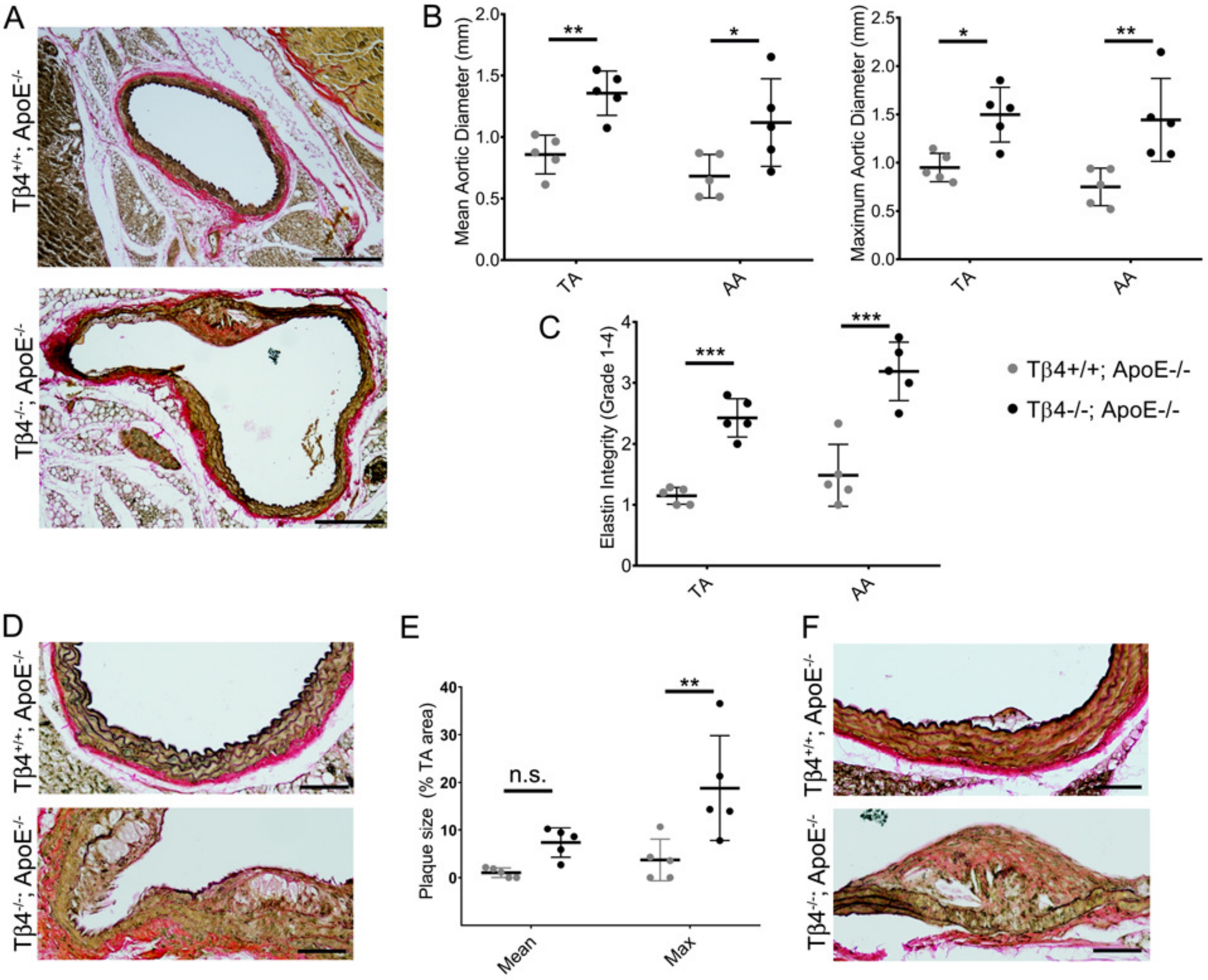


**Supplemental Figure 5. Increased predisposition to aneurysm and atherosclerotic plaque formation in regular chow-fed Tβ4$^{-/Y}$; ApoE$^{-/-}$ mice.**

Histological examination (**A**) and aortic diameter (**B**) of 8 month old Tβ4$^{+/Y}$; ApoE$^{-/-}$ and Tβ4$^{-/Y}$; ApoE$^{-/-}$ female mice, fed regular laboratory chow. Elastin integrity score shown in **C**. Incidence of plaque formation was assessed histologically (**D**) and measured in **E**. Higher magnification views of plaque to visualise composition and degeneration of underlying elastin lamellae (**F**). Data are presented as mean ± SD, with each data point representing an individual animal. Significance was calculated using one-Way ANOVA with Tukey's multiple comparison tests (**B**, **C**, **E**). n.s. = not significant; $^*p \leq 0.05$; $^{**}p \leq 0.01$; ***: $p \leq 0.001$. Scale bar: **A**: 500µm; D, F: 100µm. TA: thoracic aorta; AA: abdominal aorta.

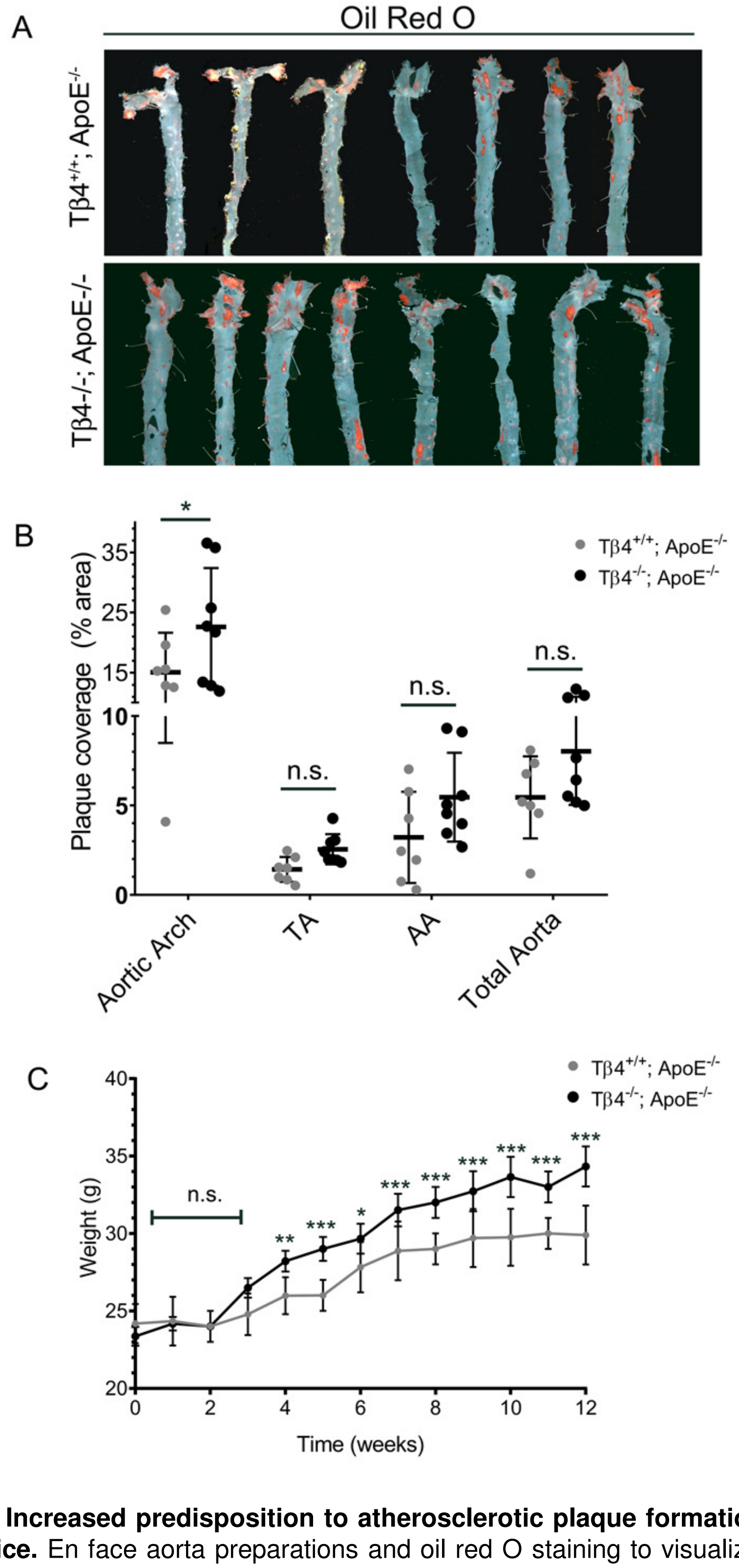


**Supplemental Figure 6. Increased predisposition to atherosclerotic plaque formation was also observed in female Tβ4$^{-/-}$; ApoE$^{-/-}$ mice.** En face aorta preparations and oil red O staining to visualize plaques in female mice fed Western diet for 12 weeks (**A**). Quantification of plaque coverage in the aortic arch, thoracic aorta (TA), abdominal aorta (AA) and total aorta (**B**). Weight gain was significantly increased in female Tβ4$^{-/-}$; ApoE$^{-/-}$, compared with Tβ4$^{-/-}$; ApoE$^{-/-}$ mice from 4-12 weeks Western diet feeding (**C**). Data are presented as mean ± SD, with each data point representing an individual animal (**B**) and n=8 (**C**). Significance was calculated using one-way ANOVA with Tukey's multiple comparison tests (**B**) and two-way ANOVA with Dunnett's post hoc tests (**C**). n.s.= not significant; *p ≤ 0.05; **p≤0.01; ***: p≤0.001.

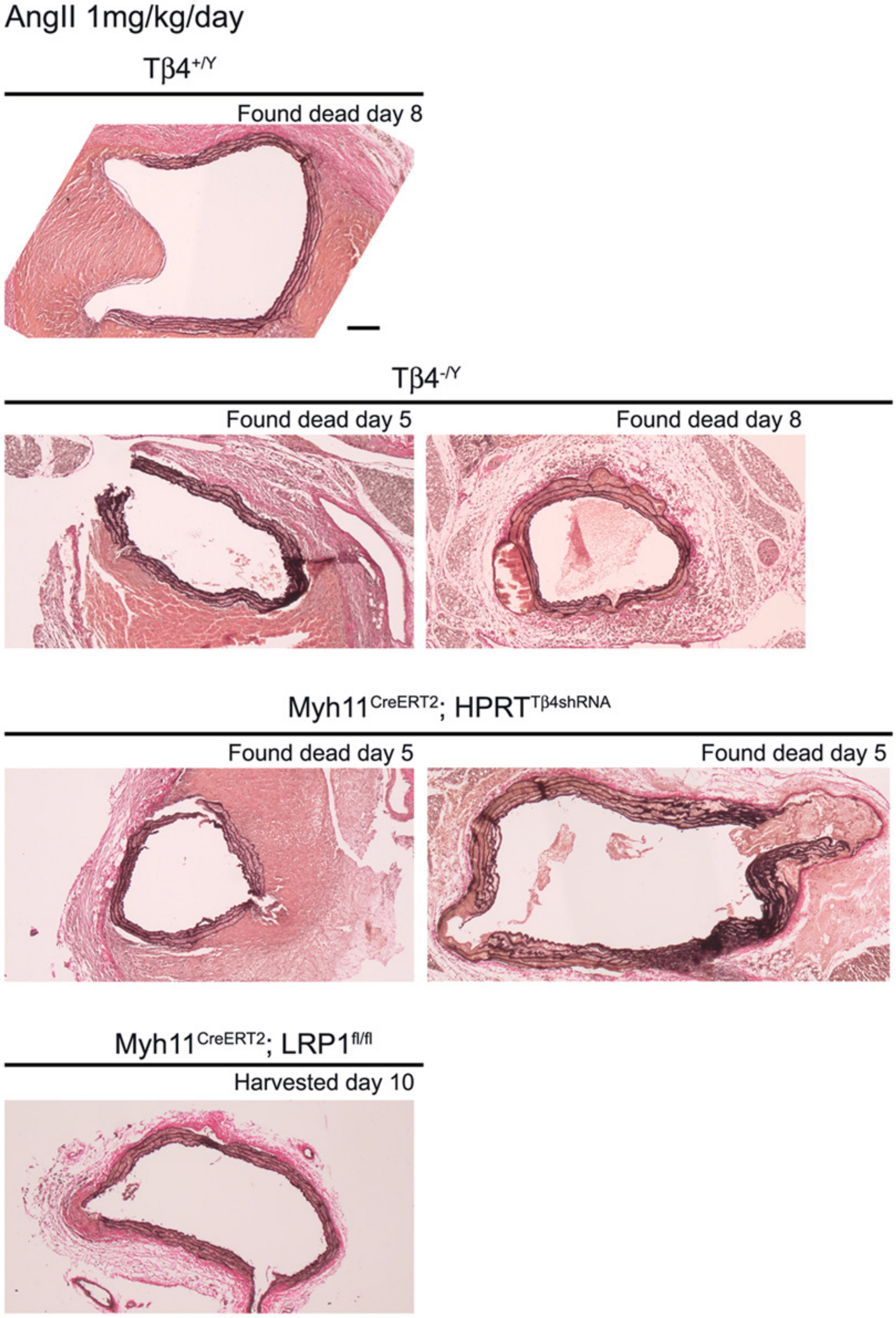


**Supplemental Figure 7. Representative examples of aortic rupture in AngII-infused mice.** A higher incidence of rupture and dissection were observed in mice lacking Tβ4 and, overall, they occurred earlier, with severe disruption of elastin lamellae and aortic dilatation. These samples could not be used for assessment of aortic diameter or elastin integrity (Figure 5) as the animals died prior to the experimental end point at day 10. Most deaths were due to aortic rupture, although occasionally dissection was detected in the same aorta, an example of which is shown for Tβ4$^{-/Y}$ (day 8). Scale bar: 200μm.

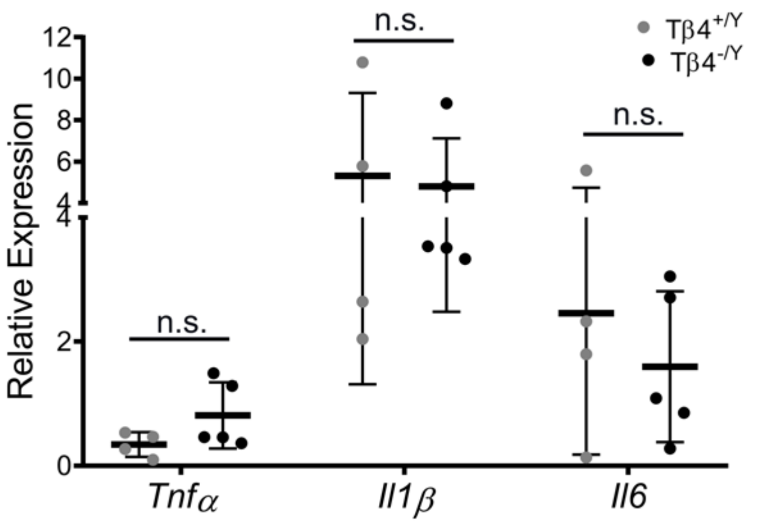


**Supplemental Figure 8. Inflammatory cytokine levels in aortic aneurysm are not significantly altered with loss of Tβ4.** qRT-PCR measurement of *Tnfα*, *Il1β* and *Il6* mRNA levels in abdominal aorta after 5 days' AngII infusion. Data are presented as mean ± SD, with each data point representing an individual animal. Significance was calculated using one-way ANOVA with Tukey's multiple comparison tests. n.s.= not significant.

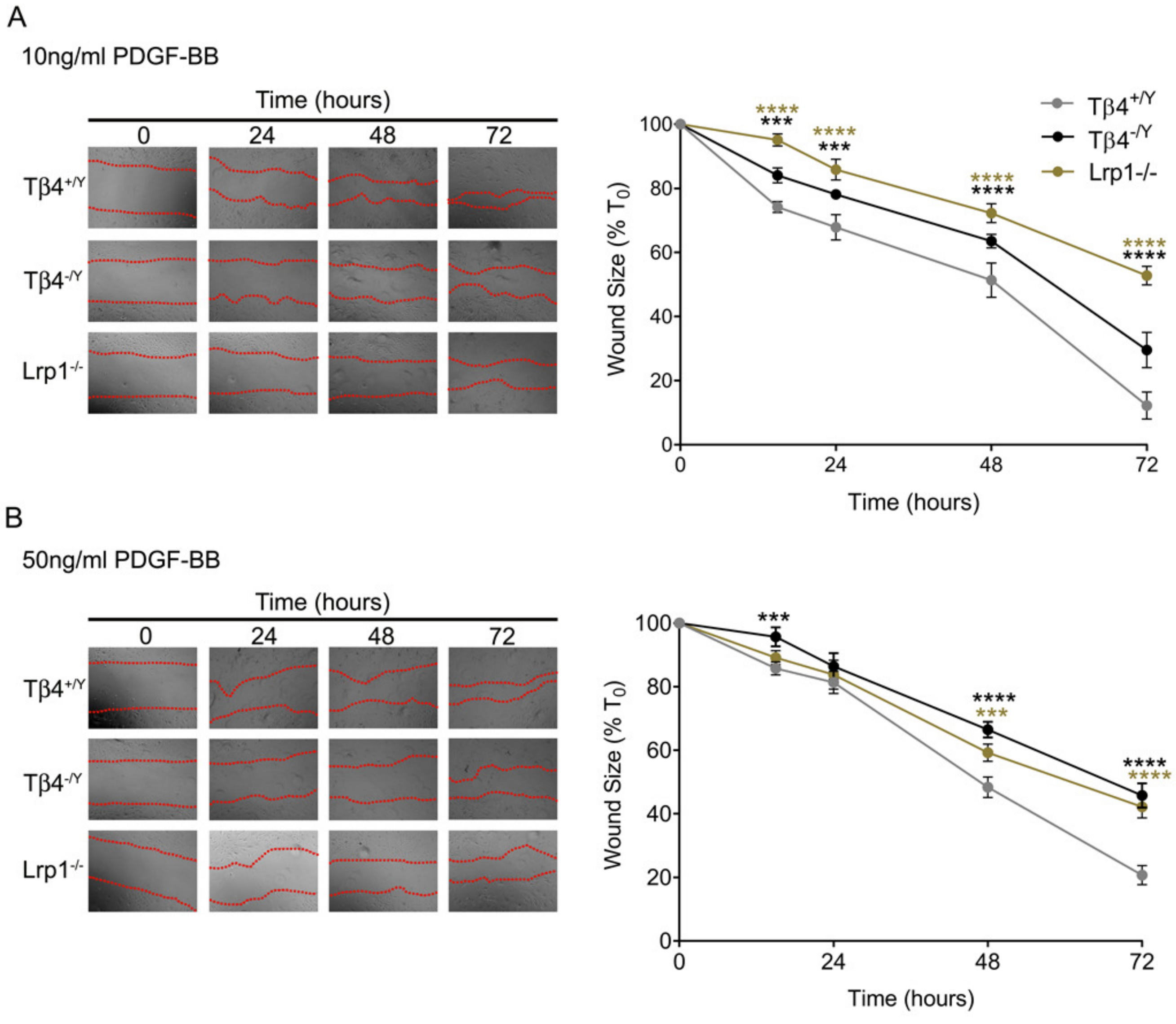


**Supplemental Figure 9. Despite augmented PDGFRβ signalling, Tβ4$^{-/Y}$ and Lrp1$^{-/-}$ aortic VSMCs migration rates are reduced, compared with control cells.**

Migration rates were determined by scratch wound assay for Tβ4$^{+/Y}$, Tβ4$^{-/Y}$ and Lrp1$^{-/-}$ aortic VSMCs, in response to 10ng/ml (**A**) or 50ng/ml (**B**) PDGF-BB. Representative images of same region of well at 0, 24, 48 and 72 hours after wounding. Data are presented as mean ± SEM from n=3 experiments. Significance was calculated using two-way ANOVA with Dunnett's post hoc tests. ***: $p \leq 0.001$; ****$p \leq 0.0001$.

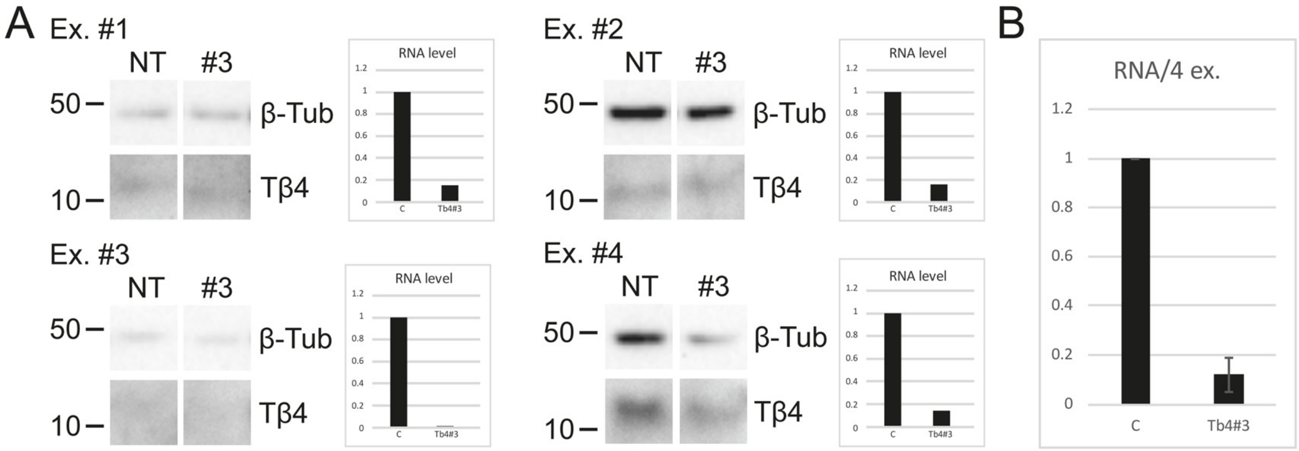


**Supplemental Figure 10. siRNA-mediated Tβ4 knockdown.** Knockdown was assessed by western blot and qRT-PCR. A minimum of 85% knockdown by qPCR was set as the criterion for inclusion in the experiments (shown in Figure 7, A-D). Knockdown data are shown for each of the individual experiments (A) and the mean qRT-PCR data (n=4).
